# GFP-Free Live-Neuron Quantitative Imaging Reveals Compartmentalization and Growth Dynamics of PolyQ Aggregates

**DOI:** 10.1101/2025.05.12.653542

**Authors:** Xiaotian Bi, Li-En Lin, Kun Miao, Lu Wei

**Author notes:** Corresponding author: Lu Wei.

## Abstract

Huntington’s Disease (HD), the most prevalent polyglutamine (polyQ) neurodegenerative disorder, features brain aggregates induced by mutant huntingtin (mHtt) proteins harboring expanded polyQ tracts. Despite extensive efforts, molecular mechanisms of polyQ aggregates remain elusive. Here we establish quantitative stimulated Raman scattering imaging of polyQ aggregates (q-aggSRS) for non-invasive investigations in live neuronal co-cultures using deuterated glutamine labeling. Q-aggSRS allows for specific visualization by targeting the distinct Raman peak from carbon–deuterium bonds, eliminating the need for bulky GFP tagging. Coupled with analysis from aggregate-tailored expansion microscopy, newly designed two-color imaging, and pulse-chase visualization, we comprehensively quantified the mHtt and non-mHtt proteins within the same aggregates across varying sizes, cell types, mHtt constructs, and subcellular locations. Our findings demonstrate a two-phase aggregate growth model with a distinct core-shell spatial organization, reveal significant heterogeneity in nucleus/cytoplasm compartmentalization specific to neurons, and identify previously unrecognized “gel-like” aggregates specifically in neuronal nuclei. These insights should advance our understanding of native polyQ aggregates and our proposed interaction coefficients may offer new quantitative parameters for developing effective HD therapies.

## Introduction

Polyglutamine (polyQ) diseases represent a group of genetic neurodegenerative disorders that cause cognitive and physical impairments, including spinocerebellar ataxias, Huntington’s Disease (HD), spinal and bulbar muscular atrophy (SMBA), and dentatorubral-pallidoluysian atrophy (DRPLA).(1-3) These diseases are characterized by the pathological expansion of CAG trinucleotide repeats, which encode polyQ tracts that exceed a critical threshold of approximately 35 repeats,(4, 5) leading to the formation of aggregates mainly in the brain. Among this family, HD is the most prevalent and widely studied, resulting from a genetic mutation in the HTT gene that induces the aggregation of mutant huntingtin (mHtt) proteins. Over the decades, extensive investigations have provided various fundamental insights into the potential mechanism of mHtt aggregation and its molecular composition.(6-9) Despite these advances, there remains no effective cure for HD. Moreover, significant phenomena, such as the predominance of nuclear aggregates in younger patients and cytoplasmic aggregates in older patients, remain poorly understood at the cellular level.(1, 8, 10) Recent research suggests that soluble mHtt oligomers may be more toxic than the aggregates, with aggregation potentially acting as a protective mechanism to sequester soluble mHtt proteins rather than directly causing cellular toxicity.(5, 9, 11-14) Nevertheless, such a mechanistic picture remains controversial, and our understanding of the structural composition and pathological roles of polyQ aggregates, particularly in the native states within neuronal environments, is limited.

Traditional biochemistry methods with *in vitro* models(15) and emerging mass spectroscopy analysis from purified aggregates offer detailed molecular views. However, these approaches involve extensive postprocessing steps such as cell lysis, centrifugation, washing, and extraction, which can introduce severe sample perturbations.(12, 16) In addition, these studies rarely employ neuronal models, relying instead on non-neuronal systems such as yeast, (14, 17-19) E. Coli,(20) or mammalian cell lines,(11, 12, 16, 21, 22) which may lead to complications in interpretation as they fail to recapitulate the neuronal environment relevant to polyQ diseases. Fluorescence microscopy allows unprecedented dynamic analysis of polyQ aggregates in live cells with high spatial resolution.(21, 23, 24) However, the common use of large fluorescent proteins such as EGFP in fusion with mHtt can disrupt its native aggregation behavior, potentially altering the original mechanism. This issue is especially critical for the widely studied disease-causing mHtt-exon1 peptide model, (25) an aberrantly spliced variant of mHtt that is less than half the size of EGFP. With the emergence of cryo-electron tomography (Cryo-ET), it is now feasible to perform ultrastructural analysis of native polyQ aggregates in unperturbed cellular environments without EGFP tagging.(26, 27) However, Cryo-ET is limited by its low-throughput, lack of quantification, and inability to capture dynamic information in live cells. An effective and high-throughput platform that allows quantitative examination of native aggregates in live and physiological-relevant models would hence be invaluable in providing new and unbiased perspectives into the previously “invisible” phenomenon of polyQ aggregates and accelerating the development of promising therapies for polyQ diseases.(2, 28-30)

Stimulated Raman scattering (SRS) microscopy, a nonlinear Raman imaging modality, has been proven to be a non-invasive optical technique for live-cell and tissue imaging with high spatial resolution.(31-34) Recently, we introduced an effective platform integrating deuterated glutamine (Gln-d_5_) labeling with SRS imaging for highly specific and quantitative analysis of polyQ aggregates in live HeLa cells, eliminating the need for EGFP tagging.(35) We demonstrated significant compositional changes caused by EGFP tagging in the live-cell context and discovered an unexpectedly high concentration of non-mHtt proteins sequestered in the aggregates.(35) However, similar to many other studies relying on non-neuronal systems, concerns arise regarding whether the observed aggregation processes fully recapitulate those occurring in live nervous systems which have a much more complex cellular environment. Not only is the intracellular environment of neurons distinct from other cell types, but their communication with glia might also play crucial roles in the formation, propagation, and potential clearance of polyQ aggregates,(36, 37) making it essential to conduct studies in a native system.

In this work, we established a quantitative SRS imaging platform of mHtt aggregates (q-aggSRS) in live primary hippocampal neuronal co-cultures through coupled AAV transduction and Gln-d_5_ metabolic incorporation to trace the aggregation formation, compartmentalization, dynamics, and composition for polyQ aggregates (Fig. 1A). By targeting the carbon-deuterium (CD) vibrational frequency in the desired cell-silent region (1800−2600 cm^-1^), SRS allowed for specific visualization of polyQ aggregates labeled by Gln-d_5_ in neuronal cells. This unique atom-labeling strategy via stable isotope replacement of carbon−hydrogen (CH) bonds with CD bonds is highly non-perturbative with minimal size difference and should largely maintain the integrity of the original hydrogen bonding network. The linear relationship between SRS signals and bond concentrations coupled with the known mHtt protein sequence uniquely allowed us to calculate absolute concentrations for both mHtt and non-mHtt proteins (c(mHtt) and c(non-mHtt)) within the same aggregates in live neurons. We comprehensively examined the influence of varied mHtt constructs, including those with and without EGFP labeling, varying polyQ tract lengths, and the deletion of the proline-rich region (Fig. 1B). Our findings also confirmed that EGFP tagging causes more severe perturbations to polyQ aggregates in neurons compared to HeLa cells.

**Fig 1.**
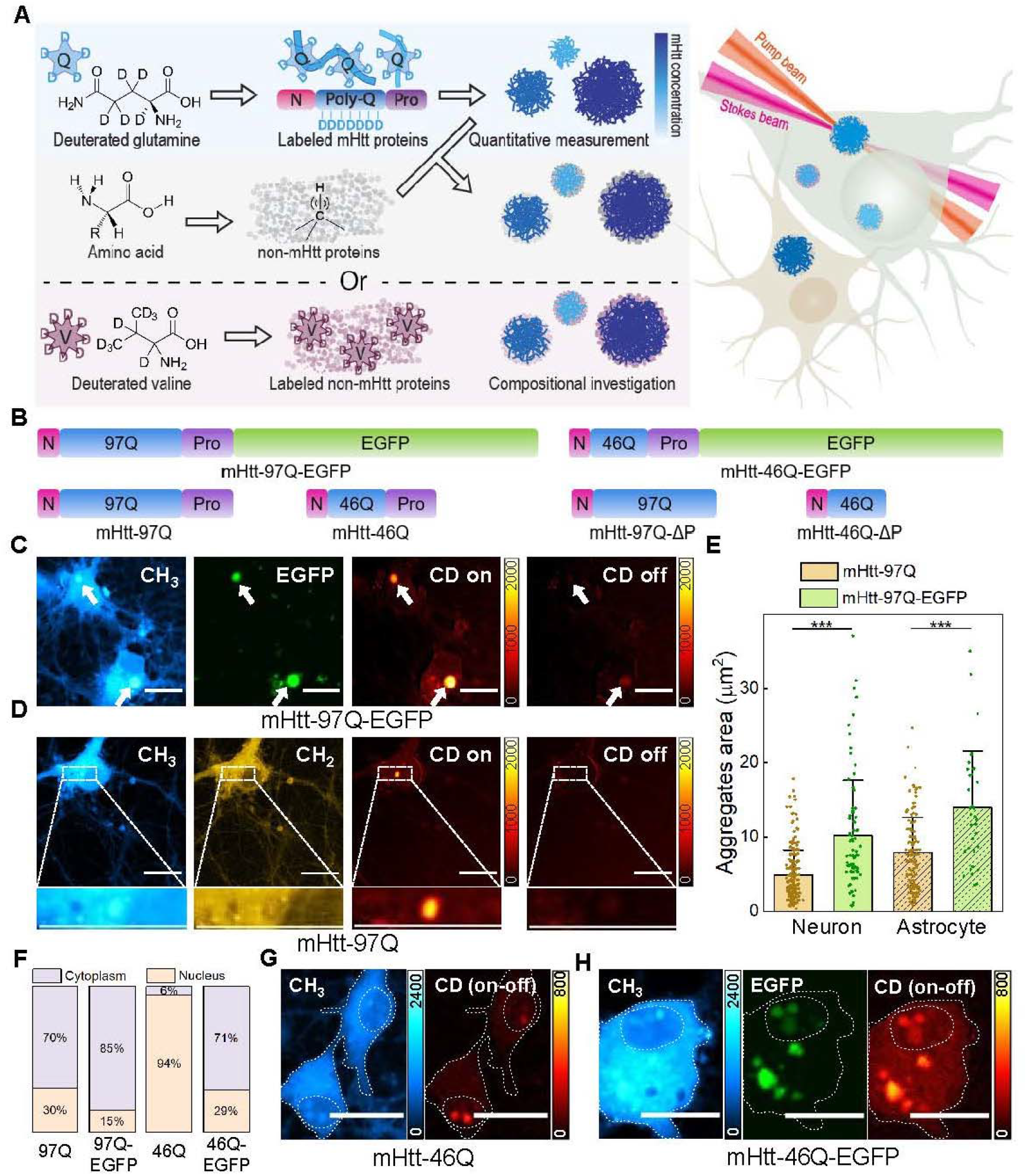
Experimental scheme for q-aggSRS imaging in hippocampal neuronal co-cultures with deuterium labeling. (a) Left: schematic illustration showing the deuterium labeling strategy for polyQ aggregates. Top: the use of deuterated glutamine (Gln-d_5_) to label mHtt proteins; middle: label-free detection of non-mHtt proteins with C-H vibrations; bottom: the use of deuterated valine (Val-d_8_) to label non-mHtt proteins. Right: Stimulated Raman Scattering (SRS) microscopy with pump and Stokes beams to image the aggregates in neuron co-cultures. (b) Plasmid constructs of various model mutant Huntingtin (mHtt) exon 1 proteins, including different polyQ lengths (46Q or 97Q), with/without the fusion of EGFP at the C terminus, and the deletion of the proline-rich region. N: N-terminal 17-amino acid fragment; 46Q/97Q: polyQ tracts with different lengths; Pro: the proline-rich region; EGFP: enhanced green fluorescent protein. (c) Fluorescence and SRS images in the same set of neurons containing mHtt-97Q-EGFP aggregates with Gln-d_5_ labeling. White arrows indicate polyQ aggregates. From left to right: the CH_3_ channel (2940 cm^-1^), EGFP fluorescence channel, CD on channel (2164 cm^-1^), and CD off channel (2036 cm^-1^). (d) SRS images in the same set of neurons containing mHtt-97Q aggregates with Gln-d_5_ labeling. The inset shows a zoomed-in view of the aggregates. From left to right: the CH_3_ channel (2940 cm^-1^), CH_2_ channel (2845 cm^-1^), CD on channel (2164 cm^-1^), and CD off channel (2036 cm^-1^). (e) Comparison of aggregate areas for mHtt-97Q and mHtt-97Q-EGFP aggregates in neurons and astrocytes, error bar: SD, *** p < 0.001. (f) Distribution of aggregates between the cytoplasm and nucleus in neurons for various mHtt constructs, including mHtt-97Q, mHtt-97Q-EGFP, mHtt-46Q, and mHtt-46Q-EGFP. (g) SRS images in the same set of neurons containing mHtt-46Q aggregates with Gln-d_5_ labeling. The dashed line outlines the cell shape and nucleus. Left: CH_3_ channel (2940 cm^-1^); right: CD on-off channel (on, 2164 cm^-1^; off, 2036 cm^-1^). (h) Fluorescence and SRS images in the same set of neurons containing mHtt-46Q-EGFP aggregates with Gln-d_5_ labeling. The dashed line outlines the cell shape and nucleus. Left: the CH_3_ channel (2940 cm^-1^); middle: EGFP fluorescence channel; right: CD on-off channel (on, 2164 cm^-1^; off, 2036 cm^-1^). Scale bars: 20 μm.

We further developed a two-color imaging strategy with co-labeling of Gln-d_5_ and deuterated valine (Val-d_8_) to unequivocally investigate mHtt and non-mHtt proteins based on the absence of valine in mHtt constructs and their distinct CD spectral features (Fig. 1A and Fig. S1). By combining expansion microscopy and live-cell pulse-chase experiments for high spatial and temporal resolution imaging, we developed a two-phase core-shell model that elucidates the spatial organization of the aggregates and illuminates a potential dynamic mechanism underlying their growth. Notably, our data offered new insights into the heterogeneous subcellular nucleus/cytoplasm compartmentalization of polyQ aggregates, phenomenon unique to neurons but not observed in astrocytes. Moreover, we identified a distinctive “gel-like” type of aggregates within neuronal nuclei that have different concentration and dynamics features compared to cytoplasmic aggerates or the previously observed “liquid-like” states.(24) All these new findings underscore the importance of using live neuronal systems with a high-throughput strategy to investigate native aggregates. Finally, we proposed an interaction coefficient (IC), defined as the concentration ratio between non-mHtt and mHtt proteins within the same aggregate to serve as a quantitative perturbation index. The IC may find utility for future drug screening needed to develop effective therapies for HD.

## Results

### The neuronal q-aggSRS imaging platform confirms severe perturbations caused by EGFP tagging

The hippocampus, a critical region involved in learning and memory, is a frequently detected site for polyQ aggregate accumulation in HD patients.(38, 39) Hippocampal atrophy and dysfunction are believed to contribute to the cognitive decline observed in the disease. Here, we adopt primary neuronal co-cultures from the rat hippocampus to obtain a stable and reproducible source of neurons for investigating polyQ aggregate formation. We utilized AAV transduction for effective gene delivery with the established disease-causing model of mHtt-exon 1, containing N-terminal 17-amino acid fragments, the polyQ tract, and the proline-rich region, which has been demonstrated to significantly contribute to mHtt-induced toxicity.(27, 40) To explore the impact of polyQ tract lengths, we selected the commonly used 97Q construct and a shorter 46Q construct, which is just above the threshold for disease initiation.(41) To validate our q-aggSRS imaging strategy in live neurons, we first benchmarked it against fluorescence imaging using mHtt-97Q-EGFP. Following AAV transduction, the medium was changed to one containing Gln-d_5_. SRS imaging targeting the CD vibration frequency at 2164 cm^-1^ (Fig. 1C, CD on) indeed clearly captured mHtt aggregates (white arrowed), consistent with that shown in fluorescence (Fig. 1C, EGFP). The much higher occurrence of glutamine in mHtt proteins compared to the general proteome provides a decent SRS signal-to-background ratio with high imaging specificity (Fig. 1C, CD on vs. CD off, 2036 cm^-1^). In parallel, SRS leveraged the established CH_3_ label-free channel (Fig. 1C, CH_3_, 2940 cm^−1^) to identify the cell morphology and map the non-labeled total proteome.

We next proceeded to image native mHtt-97Q proteins without EGFP and evaluated the impact of EGFP tagging in live neurons. While the perturbation caused by fluorescent protein tagging has been revealed by studies such as Cyro-ET and immunofluorescence, they are limited to fixed cells and purified proteins, which are subject to uncertainties from labeling specificity, antibody penetration, and protein purification length limits (25, 26, 42). Clear mHtt-97Q aggregates were detected at the CD channel (Fig. 1D, CD on, 2164 cm^-1^ vs. CD off, 2036 cm^-1^). The corresponding label-free CH_3_ and CH_2_ images provided additional information about the total proteome (Fig. 1D, CH_3_, 2940 cm^-1^) and lipidome (Fig. 1D, CH_2_, 2845 cm^-1^) distribution, with the latter indicating low lipid content within polyQ aggregates (Fig. 1D, zoomed-in view from the white-boxed area). Our results revealed that polyQ aggregates formed by mHtt-97Q were about twice as bright at the CD channel compared to those of similar sizes formed by mHtt-97Q-EGFP (Fig. 1C, D, and Fig. S2). This indicates that mHtt-97Q forms denser aggregates with higher c(mHtt) in neurons compared to those with EGFP tagging, consistent with previous findings in HeLa cells(35), and a Cryo-ET study reporting a 50% reduction in fibril density of mHtt-97Q-EGFP aggregates.(26) In addition, we observed that polyQ aggregates formed by mHtt-97Q were overall significantly smaller than those formed by mHtt-97Q-EGFP in both neurons and astrocytes (Fig. 1E and Fig. S2; see Fig. S3 and SI methods for live neuron and astrocyte identification). Interestingly, we also found that EGFP largely influenced the subcellular location of aggregates in live neurons (Fig. 1F-H). Notably, while mHtt-46Q primarily formed nuclear aggregates (Fig. 1F, G, 94% in nucleus), the EGFP-labeled counterpart dominantly formed cytoplasmic aggregates (Fig. 1F, H, 29% in nucleus). This suggests that EGFP not only alters aggregate structure but also disrupts the intracellular translocation of mHtt proteins. As a comparison, astrocytes exclusively formed cytoplasmic aggregates regardless of the mHtt construct. Our results hence highlight the importance of conducting measurements in live neurons and avoiding EGFP perturbation to accurately investigate the polyQ aggregation phenomenon.

### Q-aggSRS reveals a concentration dependence of mHtt proteins, but not non-mHtt proteins, on aggregate sizes with distinct differences in neurons and astrocytes

After establishing the q-aggSRS imaging strategy in live neuron co-cultures for studying native mHtt aggregates, we aimed to perform quantitative investigations on mHtt-97Q with high spatial resolution (Fig. 2A-D). We observed polyQ aggregates of varying sizes in the cytoplasm (Fig. 2A, arrowed) that correspond to drastically different calculated c(mHtt) (Fig. 2C, arrowed, see SI methods and Fig. S4 for calculation details) and CH/CD ratios (Fig. 2D, arrowed), directly measured through dividing the aggregate signals at the CH_3_ channel (Fig. 2B, arrowed) by those at the CD channel (Fig. 2A, arrowed). CH/CD ratios serve as a surrogate indicator for the ratios of non-mHtt/mHtt proteins, since CD signals from Gln-d_5_ mainly originate from mHtt proteins, while the CH_3_ signals from non-Gln amino acids (a.a.), mainly represent the sequestered non-mHtt proteins within the aggregates. With CH/CD ratios and the precise sequence of mHtt-exon1, absolute c(mHtt) and c(non-mHtt) within the same aggregates could be calculated correspondingly (SI methods). The c(non-mHtt) is quantified using the average proteome protein size as a reference scale.(43, 44) Compared to the fibril density measurement by cyro-ET,(26) our absolute protein concentrations should provide a complementary quantitative assessment that reflects the local density of protein monomers.

**Fig 2.**
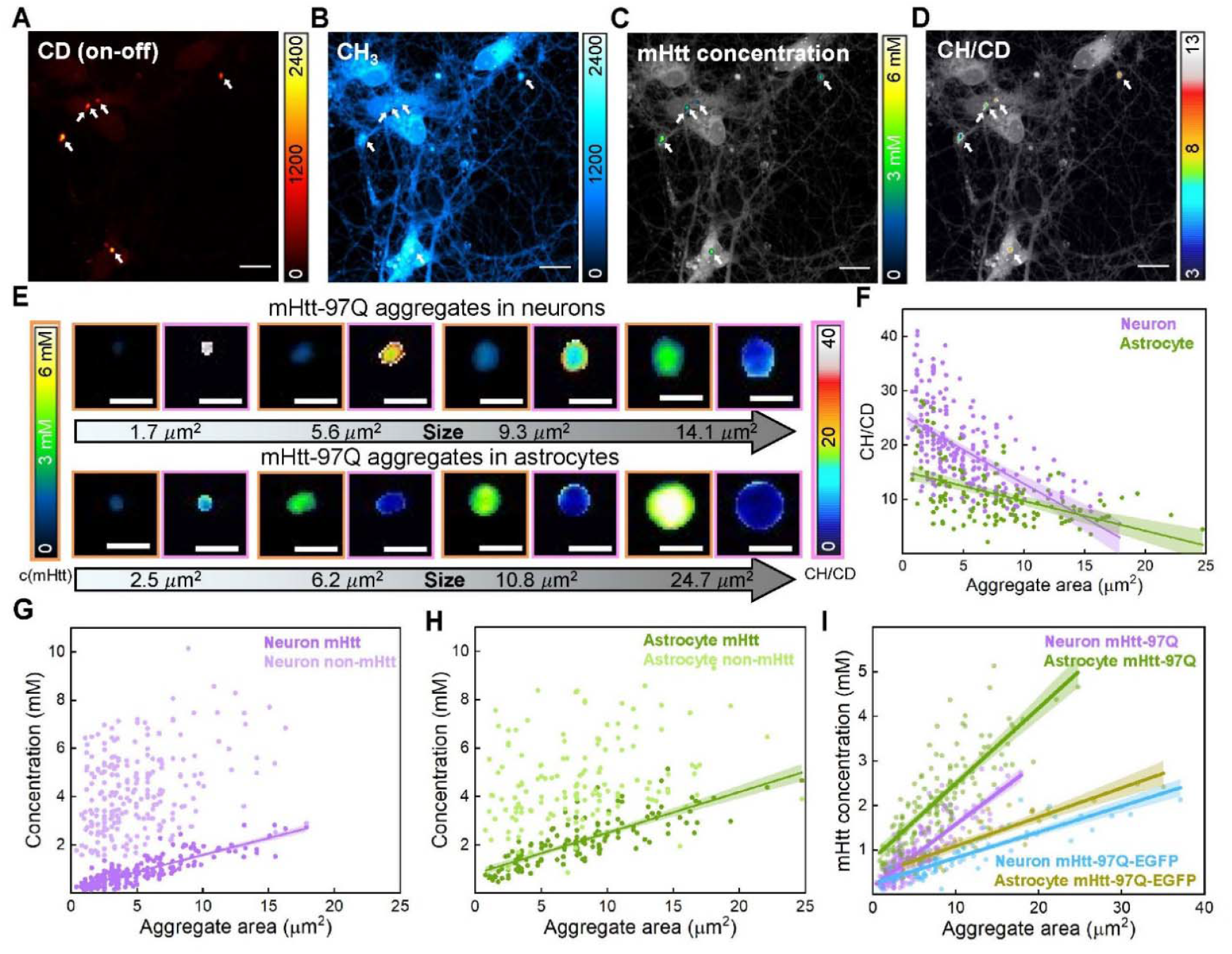
Quantitative measurement of mHtt-97Q cytoplasmic aggregates with q-aggSRS. (A-B) SRS image at the CD on-off channel (A, on, 2164 cm^-1^; off, 2036 cm^-1^) and the CH_3_ channel (B, 2940 cm^-1^) in the same set of neurons containing mHtt-97Q aggregates with Gln-d_5_ labeling. (C-D) Image displaying the calculated c(mHtt) (C) and CH/CD ratios (D) within aggregates with the grey color representing the CH_3_ channel (2940 cm^-1^) in the same set of neurons as (A). Scale bar: 20 μm, white arrows indicate aggregates of different sizes. (E) Representative aggregates with increasing sizes, top: in neurons; bottom: in astrocytes; left: c(mHtt) image with orange border; right: CH/CD ratiometric image with pink border, scale bar: 5 μm. (F) Scatter plots with linear fittings between CH/CD ratios and aggregate areas of polyQ aggregates in neurons (purple, *y* = -1.25*x* + 25.42, Pearson’s *r* = -0.56) and astrocytes (green, *y* = -0.55*x* + 15.22, Pearson’s *r* = -0.50). (G) Scatter plots with linear fittings of c(mHtt) (dark purple, *y* = 0.14*x* + 0.18, Pearson’s *r* = 0.84) and c(non-mHtt) (light purple, Pearson’s *r* = 0.35) over aggregate areas of polyQ aggregates in neurons. (H) Scatter plots with linear fittings of c(mHtt) (dark green, *y* = 0.17*x* + 0.81, Pearson’s *r* = 0.81) and c(non-mHtt) (light green, Pearson’s *r* = 0.28) over aggregate areas of polyQ aggregates in astrocytes. (I) Scatter plots with linear fittings of c(mHtt) over aggregate areas of polyQ aggregates (mHtt-97Q/ mHtt-97Q-EGFP) in neurons and astrocytes. mHtt-97Q-EGFP: neuron c(mHtt), cyan blue: *y* = 0.058*x* + 0.25, Pearson’s *r* = 0.88; astrocyte c(mHtt), mustard yellow: *y* = 0.065*x* + 0.44, Pearson’s *r* = 0.88. (F-I) A 95% confidence band is plotted with the fitted line.

Zoomed-in views illustrate aggregates of increasing sizes (Fig. 2E, left panels with orange borders: c(mHtt); right panels with pink borders: CH/CD) in both neurons (Fig. 2E, top row) and astrocytes (Fig. 2E, bottom row). Comprehensive analysis of hundreds of aggregates across different sizes reveals that CH/CD ratios (Fig. 2F) exhibit a weak negative correlation with aggregate sizes, which originates from a strong positive correlation between calculated c(mHtt) and aggregate sizes (Pearson’s r >0.8) in both neurons (Fig. 2G, dark purple) and astrocytes (Fig. 2H, dark green). In contrast, c(non-mHtt) sequestered in the same aggregates is intriguingly high, reaching up to 10 mM, but shows a much weaker and scattered dependence on aggregate sizes (Pearson’s r <0.4) (Fig. 2G, light purple, and Fig. 2H, light green). This indicates that the amount of trapped non-mHtt proteins is independent of aggregate size and the protein entrapment mechanisms may be different for mHtt and non-mHtt proteins. To our knowledge, this is the first time that quantitative mapping of polyQ aggregates has been achieved in live neurons and astrocytes.

We further found that aggregates of the same sizes consistently present a 2-fold higher c(mHtt) in astrocytes (Fig. 2I, dark green) compared to those in neurons (Fig. 2I, dark purple). Interestingly, the fitting curves showed lower intercepts in neurons compared to astrocytes (Fig. 2I, dark purple vs. dark green, and Fig. S5A, B), which implies that neurons may have a lower concentration threshold for aggregate formation, and are likely more susceptible to perturbations of mHtt proteins.(36, 37, 45) Such heterogeneity between neurons and astrocytes became much smaller for the mHtt-97Q-EGFP aggregates (Fig. 2I, cyan blue vs. mustard yellow), again emphasizing the influence of EGFP labeling in a direct live-cell context. Additionally, aggregates were found to appear in both the cell body and axons/dendrites regions. While the c(mHtt) vs. aggregate size relationship remains the same across different subcellular locations (Fig. S6A), the CH/CD ratios in the cell body were slightly higher (Fig. S6B), likely reflecting differences in the local subcellular environment. However, due to the similar quantification matrix, we will focus on the analysis of aggregates in the cell bodies.

### Expansion and two-color q-aggSRS imaging reveal a core-shell aggregate structure

As shown in Fig. 2E, the CH/CD ratiometric images (right panels with pink borders) imply a surprising core-shell structure, where mHtt proteins are concentrated in the core of aggregates and non-mHtt proteins are relatively more prominent at the periphery. We hence sought to confirm and understand such a core-shell spatial distribution of mHtt and non-mHtt proteins within the same aggregates with enhanced spatial and spectral resolution. First, we applied Raman-tailored sample-expansion microscopy to perform q-aggSRS VISTA (Vibrational Imaging of Swelled Tissues and Analysis),(46) a super-resolution Raman imaging technique previously developed in our laboratory. VISTA has been applied to probe both polyQ and amyloid beta protein aggregates in cells and tissues with validated isotropic expansion(47). Neurons containing mHtt-97Q aggregates were embedded in polymer hydrogels and then expanded isotropically by 2-fold to reach an effective resolution below 200 nm. Indeed, the overlay image (Fig. 3A, right) of the CH_3_ (Fig. 3A, left) and the CD (Fig. 3A, middle) channel clearly confirms a core-shell structure with non-mHtt proteins being relatively more abundant in the shell.

**Fig 3.**
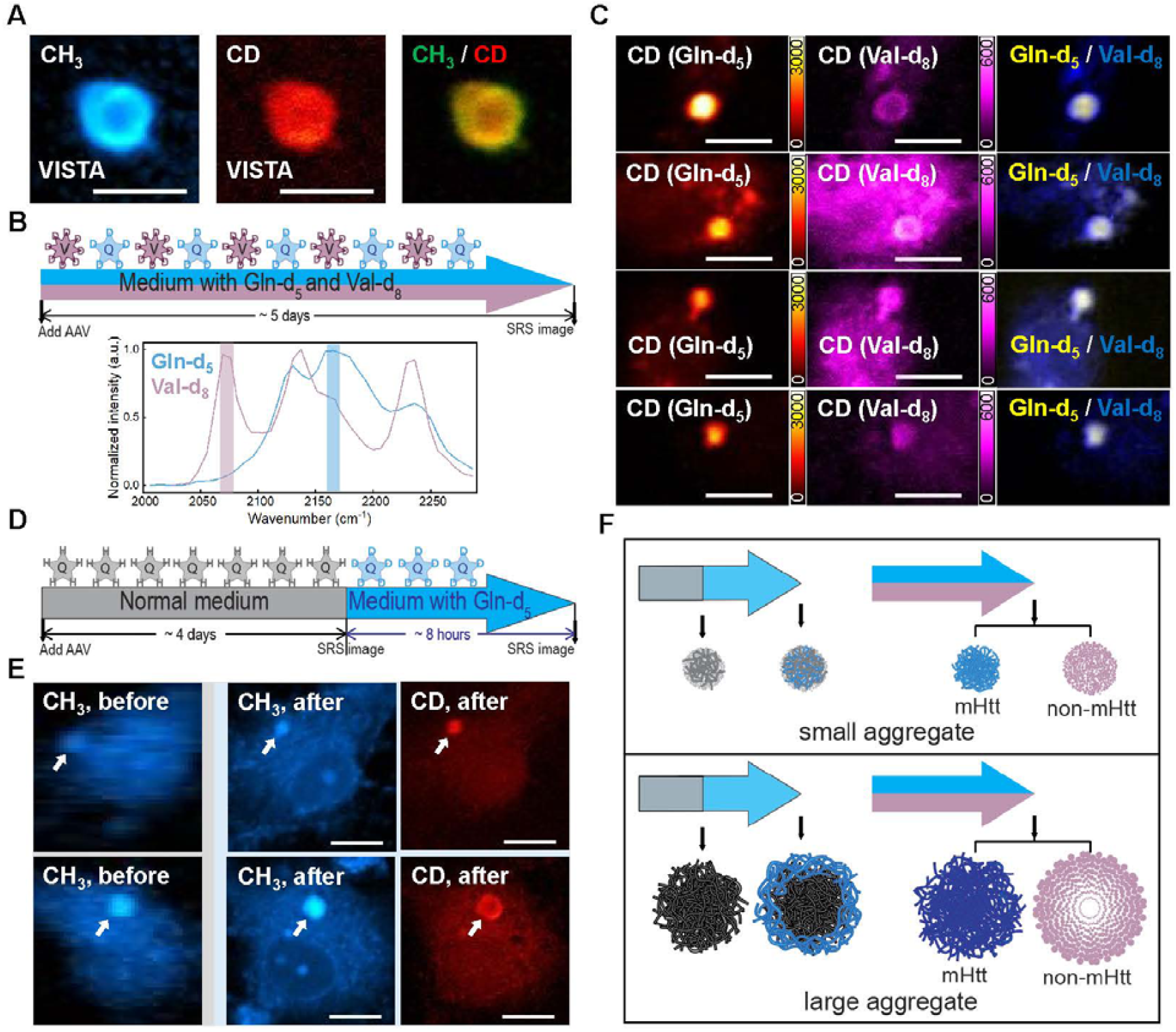
Structural investigation and dynamic tracking of mHtt-97Q cytoplasmic aggregates. (A) Q-aggSRS images from the VISTA sample (2-fold expansion). From left to right: the CH_3_ channel (2940 cm^-1^), the CD channel (2164 cm^-1^), merged image of the CH_3_ and CD channels, scale bar: 10 μm. (B) Top: Experimental design of two-color imaging by co-incubation of Gln-d_5_ and Val-d_8_. Bottom: SRS spectra of polyQ aggregates with either Gln-d_5_ (blue) or Val-d_8_ (purple) incorporation. The two colored bands indicate SRS acquisition positions (2068 cm^-1^, 2164 cm^-1^) for unmixing. (C) Two-color imaging of Gln-d_5_ labeled mHtt proteins and Val-d_8_ labeled non-mHtt proteins for 4 sets of aggregates, scale bar: 10 μm. (D) Experimental design for the pulse-chase labeling experiment. (E) Q-aggSRS images at the CH_3_ channel (2940 cm^-1^) and CD channel (2164 cm^-1^) in the same set of neurons before and after the medium replacement with white arrows indicating polyQ aggregates. The top row shows a diffused CD signal across the aggregate, while the bottom row shows a shell structure of the CD signal in the aggregate, scale bar: 10 μm. (F) Proposed two-phase aggregate growth model for small aggregates (top) and large aggregates (bottom), building upon the pulse-chase experiment (left) and the spatial distribution of mHtt proteins and non-mHtt proteins within the same polyQ aggregate (right).

Second, we devised a new two-color q-aggSRS strategy (Fig. 3B, top) by co-labeling live neurons with Gln-d_5_ and deuterated valine (Val-d_8_), harnessing differences in Raman spectral features arising from their distinct chemical structures (Fig. 3B, bottom). Since valine is absent in mHtt protein sequences, all Val-d_8_ signals originate exclusively from newly synthesized non-mHtt proteins, eliminating any crosstalk contributed by mHtt proteins, which exist in the CH_3_ signals. Such a two-color strategy more clearly resolved the core-shell analysis. By incubating cells with Gln-d_5_ and Val-d_8_ substituted culture medium and performing two-color q-aggSRS imaging (Fig. 3B, and see SI methods for unmixing details), our results consistently confirmed clear core-shell structures (Fig. 3C, right, merged), with non-mHtt proteins (Fig. 3C, middle, Val-d_8_) more enriched in the shell and mHtt in the core (Fig. 3C, left, Gln-d_5_). This spatial organization becomes more pronounced as aggregate sizes increase (Fig. 3C, top two rows).

### Pulse-chase q-aggSRS reveals two-phase growth dynamics

One unique advantage of our metabolic labeling strategy is the ability to distinguish newly synthesized proteins from the pre-existing pool by a pulse-chase medium change over time. To investigate the growth dynamic of cytoplasmic mHtt-97Q aggregates in live neurons, we designed a pulse-chase time-lapse experiment (Fig. 3D). Neurons were initially cultured in a regular medium for 4 days post-AAV addition to allow sufficient protein expression, followed by first round of SRS imaging at the CH_3_ channel (Fig. 3E, CH_3_, before) to confirm the formation of polyQ aggregates (arrowed). Here, we chose very low laser power and rapid acquisition time to minimize potential photodamage to neurons (Fig. 3E, CH_3_, before). The regular medium was then replaced with the Gln-d_5_ medium for 8 hours (Fig. 3D), after which the same neurons were re-imaged at both the CH_3_ and the CD channels (Fig. 3E, CH_3_, after and CD, after). Here, the CD signals indicate newly synthesized mHtt proteins incorporated into the same aggregates after the medium change. Interestingly, representative q-aggSRS images revealed two distinct patterns of CD signal distribution: a uniform distribution throughout smaller aggregates (Fig. 3E, top row, arrowed) and an exclusive localization to the periphery in larger aggregates (Fig. 3E, bottom row, arrowed). This result suggests that large aggregates represent a more compact and inert state as newly sequestered mHtt proteins are mostly restricted to the outer layer of large aggregates, whereas small aggregates allow newly sequestered mHtt proteins to diffuse throughout the core, implying a more flexible structural organization.

Collectively, we propose a two-phase composition and growth model between the small and large polyQ aggregates (Fig. 3F). According to Cryo-ET high-resolution structural studies, mHtt forms polyQ aggregates in the form of fibrils.(27, 48, 49) Based on this, we propose that small aggregates have a relatively loose fibril structure with higher flexibility and diffusion rate, hence allowing newly sequestered proteins to freely diffuse into the core (Fig. 3F, top row, left). In addition, they form a weak core-shell compositional structure featuring a minor exclusion of non-mHtt proteins in the center (Fig. 3F, top row, right). In large aggregates with higher c(mHtt), the core is denser and more compact, preventing further diffusion of newly formed mHtt fibrils into the core and leading to a shell-like distribution of newly sequestered mHtt surrounding the dense core (Fig. 3F, bottom row, left). This effect is likely more pronounced for non-mHtt proteins, hence resulting in an obvious core-shell compositional structure between mHtt and non-mHtt cytosolic proteins in large aggregates (Fig. 3F, bottom row, right). Our core-shell model provides new insights and aids in the understanding of polyQ aggregate formation, growth, and structural composition.

### Nuclear polyQ aggregates are dominant for the shorter Q length and feature lower c(mHtt) than cytoplasmic aggregates

The above discussions feature cytoplasmic polyQ aggregates, which are the major form in mHtt-97Q (Fig. 1F, 70% in cytoplasm). For mHtt-46Q with shorter Q length, nuclear aggregates became dominant (Fig. 1F, 94% in nucleus, Fig. 4A). We confirmed that the observed aggregates were not nucleoli, which would appear more prominent in the CH_3_ channel (Fig. 4A). We further found that mHtt-46Q aggregates exclusively localized in the cytoplasm of astrocytes (Fig. 4A, 4B), revealing stark differences in aggregation compartmentalization between neurons and astrocytes. The c(mHtt) of nuclear aggregates was also found to be significantly lower and the CH/CD ratios were significantly higher compared to cytoplasmic ones (Fig. 4C, 4D). This suggests a more preferential sequestration of non-mHtt proteins in nuclear aggregates (Fig. S7), likely contributing to higher toxicity as one suggested toxicity mechanism involves the depletion of functional proteins by aggregate formation(4, 16, 18, 50-53). In addition, we found that while there was typically one cytoplasmic aggregate per cell (Fig. 4E, green and purple histograms with distribution curves), there were usually multiple nuclear aggregates per cell (Fig. 4E, light blue histogram with distribution curve). These findings indicate distinct features between nuclear and cytoplasmic aggregates,(42) emphasizing the role of the subcellular environment in influencing aggregate formation.

**Fig 4.**
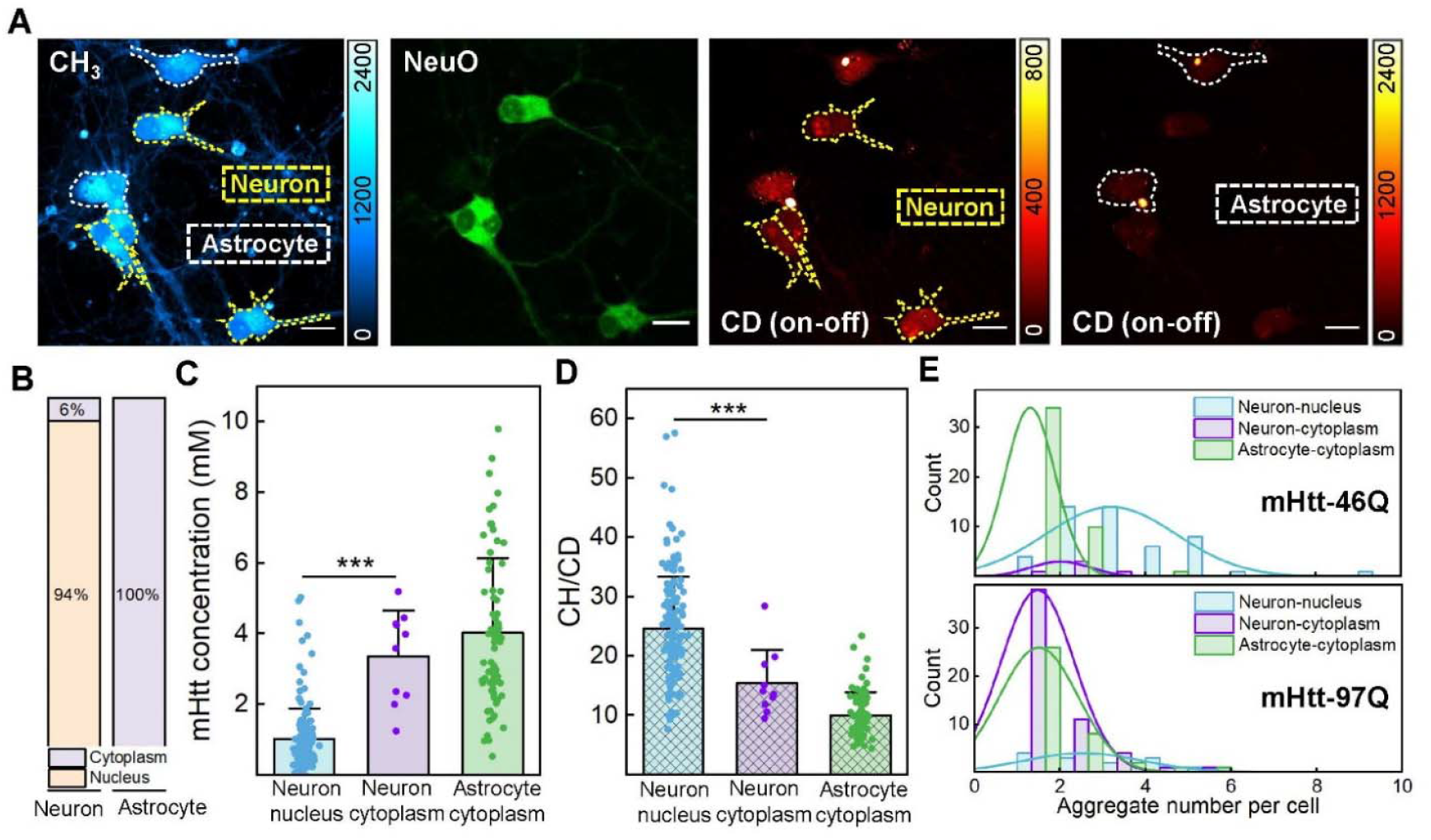
Quantitative measurement of mHtt-46Q nuclear aggregates with q-aggSRS. (A) Fluorescence and SRS images in the same set of neuronal co-cultures containing mHtt-46Q aggregates. From left to right: the CH_3_ channel (2164 cm^-1^) with the outline for neurons (dashed, yellow boxes) and astrocytes (dashed, white boxes), the fluorescence image showing NeuO(61) staining for neurons, the CD on-off channel (on, 2164 cm^-1^; off, 2036 cm^-1^) image showing nuclear aggregates in neurons, the CD on-off channel (on, 2164 cm^-1^; off, 2036 cm^-1^) image showing cytoplasmic aggregates in astrocytes under different contrast, scale bar: 20 μm. (B) Distribution of nuclear and cytoplasmic aggregates for mHtt-46Q in neurons and astrocytes. (C-D) c(mHtt) (C) and CH/CD ratios (D) of mHtt-46Q aggregates in different subcellular compartments, including neuron nucleus, neuron cytoplasm, and astrocyte cytoplasm, *** p < 0.001. (E) Histogram plots showing the distribution of aggregate numbers per cell for mHtt-46Q and mHtt-97Q in different subcellular compartments, including neuron nucleus, neuron cytoplasm, and astrocyte cytoplasm.

### The deletion of the proline-rich region significantly changes c(non-mHtt) and aggregate structures

We next sought to examine the effect of deleting the proline-rich region in mHtt constructs, using mHtt-46Q-ΔP and mHtt-97Q-ΔP. Previous yeast survival studies showed that deleting the proline-rich region enhances the toxicity of mHtt.(17, 54) However, the relationship between mHtt sequences and their toxicity remains unclear. Our experiments revealed that the deletion of the proline-rich region slightly increased the percentage of nuclear aggregates (Fig. 5A vs. Fig. 1F). Representative q-aggSRS images at both the CD and CH_3_ channels of mHtt-46Q-ΔP aggregates in a neuron show a much higher contrast for aggregates at the CH_3_ channel (Fig. 5B), compared to mHtt-46Q aggregates (Fig. 4A, CH_3_), indicating a much-increased sequestration of non-mHtt proteins in the mHtt-46Q-ΔP aggregates. Subsequent statistical analysis indeed confirmed the obvious increase in c(non-mHtt) for both mHtt-46Q-ΔP (Fig. 5C) and mHtt-97Q-ΔP (Fig. 5D) across different subcellular compartments in neurons and astrocytes, with only minor changes for c(mHtt) (Figs. S8A, B). This finding is again consistent with the proposed toxicity mechanism of aggregates, which involves the depletion of essential structural (i.e. actins) and functional (i.e. chaperones) cytosolic proteins.(4, 16, 18, 50-53) The observed increase in absolute c(non-mHtt) may provide quantitative support for the enhanced toxicity associated with the deletion of the proline-rich region.

**Fig 5.**
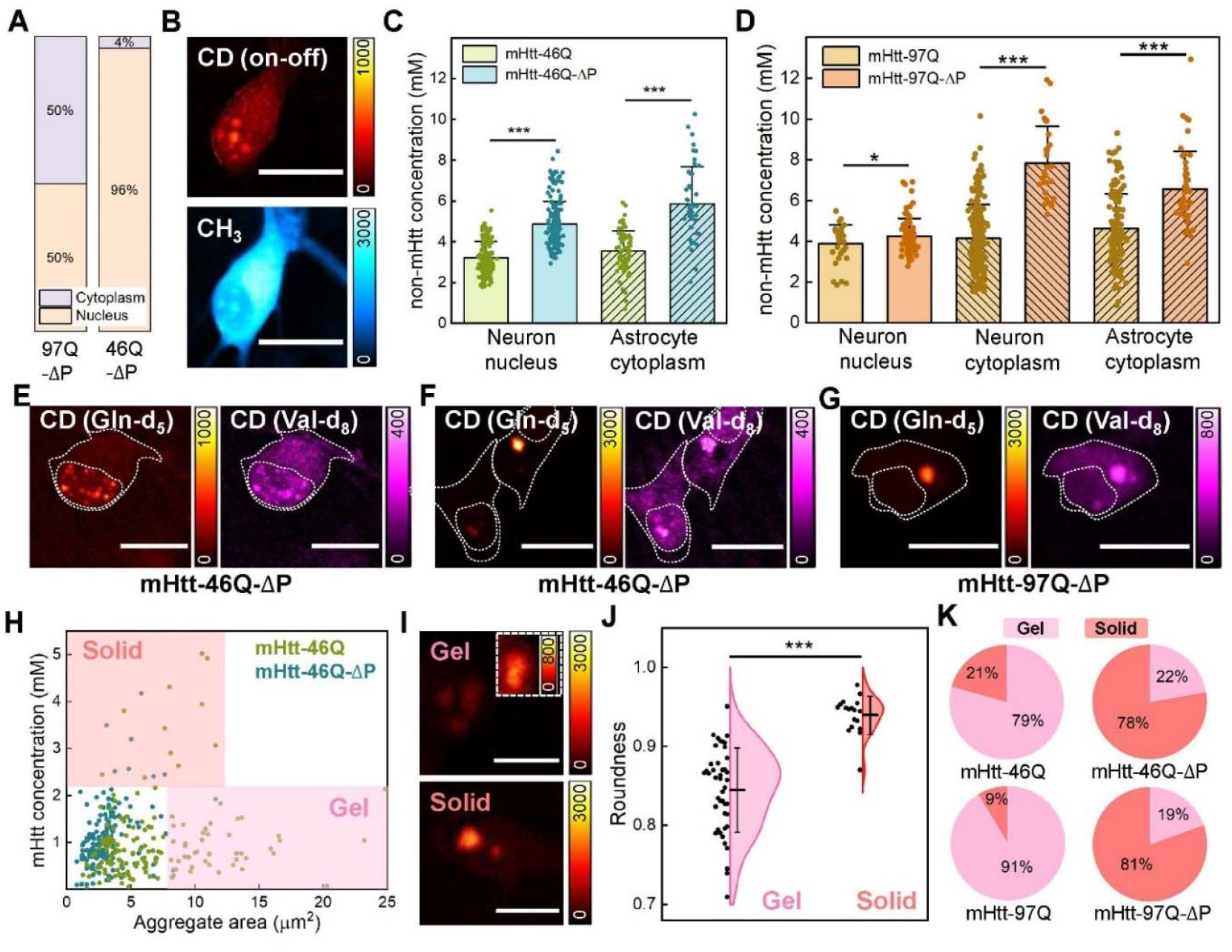
Quantitative measurement of mHtt-46Q-ΔP and mHtt-97Q-ΔP aggregates with q-aggSRS. (A) Distribution of nuclear and cytoplasmic aggregates in neurons for mHtt-46Q-ΔP and mHtt-97Q-ΔP. (B) q-aggSRS images at the CD (top, 2164 cm^-1^) and CH_3_ (bottom, 2940 cm^-1^) channels in the same set of neuron containing mHtt-46Q-ΔP aggregates with Gln-d_5_ labeling, scale bar: 20 μm. (C-D) Comparison of c(non-mHtt) in different subcellular compartments, including neuron nuclei, neuron cytoplasm, and astrocyte cytoplasm, for mHtt-46Q/mHtt-46Q-ΔP (C) and mHtt-97Q/mHtt-97Q-ΔP (D), * p < 0.05, *** p < 0.001. No neuronal cytoplasm data is shown in (C) due to an insufficient number of aggregates detected, as the majority of aggregates are in neuronal nuclei. (E-G) Two-color imaging of Gln-d_5_ labeled mHtt proteins and Val-d_8_ labeled non-mHtt proteins for mHtt-46Q-ΔP nuclear aggregates (E), mHtt-46Q-ΔP cytoplasmic aggregates (F), mHtt-97Q-ΔP cytoplasmic aggregates (G). The dashed lines outline the cell shape and nucleus, scale bar: 20 μm. (H) Scatter plot of c(mHtt) vs nuclear aggregate areas for mHtt-46Q and mHtt-46Q-ΔP, indicating a bifurcation of concentration-area dependence. The pink region (lower right quadrant) indicates the nuclear gel-like aggregates, which feature low concentration-area dependence; the coral region (upper left quadrant) indicates the nuclear solid-like aggregates, with high concentration-area dependence. (I) Q-aggSRS images of representative nuclear gel-like aggregates (top) and solid-like aggregates (bottom) at the CD on-off channel (on, 2164 cm^-1^; off, 2036 cm^-1^), scale bar: 10 μm. The white box inset shows the same gel-like aggregate but with enhanced contrast. (J) Violin plot comparing the roundness of nuclear gel-like and solid-like mHtt aggregates, *** p < 0.001. (K) Pie charts showing the proportion of gel-like and solid-like nuclear aggregates across mHtt constructs of mHtt-46Q, mHtt-46Q-ΔP, mHtt-97Q, and mHtt-97Q-ΔP.

Interestingly, our two-color imaging through the co-incubation of Gln-d_5_ and Val-d_8_ demonstrated that ΔP sequences always resulted in a homogenous aggregate structure between mHtt and non-mHtt proteins in both nucleus and cytoplasm, regardless of aggregation sizes and Q length (Fig. 5E-G). This drastically differs from the previously characterized core-shell structure observed from mHtt-97Q, even for large cytoplasmic aggregates formed by mHtt-97Q-ΔP (Fig. 5G). Previous protein structural studies identified the proline-rich region as a helix breaker that disrupts the helical structure formed by polyQ tracts. For ΔP sequences, it is plausible that the helical structures remain relatively intact and hence promote mHtt aggregation.(55-58) The increased aggregation capability in ΔP sequences might also strengthen the interaction between mHtt and non-mHtt proteins and cause more effective sequestration of non-mHtt proteins to the core regardless of aggregate sizes, highlighting the importance of further understanding of the interaction between sequestered mHtt and non-mHtt proteins.

### An unrecognized “gel-like” state is revealed for nuclear aggregates

Similar to our analysis for cytoplasmic aggregates, we next plotted the c(mHtt) dependence on the nuclear aggregate sizes across all four sequences: mHtt-46Q and mHtt-46Q-ΔP (Fig. 5H), and mHtt-97Q and mHtt-97Q-ΔP (Fig. S9A). Surprisingly, we observed a drastic bifurcation, revealing two distinct populations, highlighted in separate quadrants (Fig. 5H and S9A). The upper-left population (Fig. 5H, shaded coral) has a strong positive correlation between c(mHtt) and aggregate sizes, similar to the trend shown above in cytoplasmic aggregates. We hence defined this population as the “solid-like” aggregates, featuring dense mHtt core with lower mobility. In contrast, the lower-right population (Fig. 5H, shaded pink) shows no correlation between c(mHtt) and aggregate sizes. Since their density remains uncorrelated with variations in size, but their protein mobility is lower compared to the known “liquid” form, (24) we defined this low-concentration type of aggregates as “gel-like”, close to the semi-solid properties of a gel. Representative images show that the “gel-like” aggregates appear elongated (Fig. 5I, top, contrast-adjusted image shown in the while-dashed box), whereas the “solid-like” aggregates are more circular with significantly higher CD intensities (Fig. 5I, bottom). We next calculated the roundness, defined as a ratio of 4*π*\*area divided by the square of the perimeter, where a value closer to 1 indicates a shape closer to a perfect circle. Statistical analysis confirmed a significant difference in roundness between the two types, with “solid-like” aggregates being more circular (Fig. 5J). Based on the roundness, we again note that this “gel-like” state is distinct from previously reported “liquid-like” state which exhibits a more circular shape due to its fluidity.(24) These “gel-like” aggregates we discovered for the first time, defined based on their physical properties of concentration, fluidity, and morphology, are expected have higher diffusion flexibility for sequestration compared to “solid-like” ones, but possess less fluidity compared to the “liquid-like” state. Interestingly, we found that the percentage of “gel-like” nuclear aggregates largely decreases for both mHtt-46Q-ΔP and mHtt-97Q-ΔP compared to their counterparts (Fig. 5K). Along with the above two-color imaging results (Fig. 5E-G), this again indicates that the deletion of the proline-rich region should promote the binding interactions between sequestered mHtt and non-mHtt proteins, further changing the properties of polyQ aggregates.

## Discussion

We established the neuronal q-aggSRS microscopy platform for live hippocampal neuron co-cultures and conducted a comprehensive and quantitative analysis of mHtt polyQ aggregates under various conditions in an EGFP-free manner. Adding more evidence to previous studies with fixed-cells and purified proteins, we substantiated the increased perturbation of EGFP tagging for interrogating the aggregate properties in live-cell context with high specificity and more pathologically relevant models. We calculated the concentrations ofr mHtt and non-mHtt proteins and revealed varied concentration dependence on sizes. Utilizing expansion q-aggSRS imaging, along with two-color and pulse-chase time-lapse imaging, our data established a two-phase growth model with detailed compositional structures for small and large polyQ aggregates *in situ* from live neurons. Interestingly, we revealed notable heterogeneity in subcellular location for nuclear and cytoplasmic aggregates across different mHtt constructs and between neurons and astrocytes. Specifically, we only detected cytoplasmic aggregates in astrocytes, while both nuclear and cytoplasmic aggregates were found to exist in neurons (Fig. 6A). This highlights the importance of using neurons for polyQ studies, as other cell models may not fully replicate the native states and subcellular environment. In neuronal nuclei, we identified unique “gel-like” aggregates, which have c(mHtt) independent of aggregate sizes and feature a homogeneous spatial distribution of mHtt and non-mHtt proteins (Fig. 6B, pink, and S9C). This is drastically different from cytoplasmic aggregates, which have a strong linear correlation between c(mHtt) and aggregate sizes and show a relatively homogeneous composition in small aggregates but a clear core-shell structure in large aggregates (Fig. 6B, coral).

**Fig 6.**
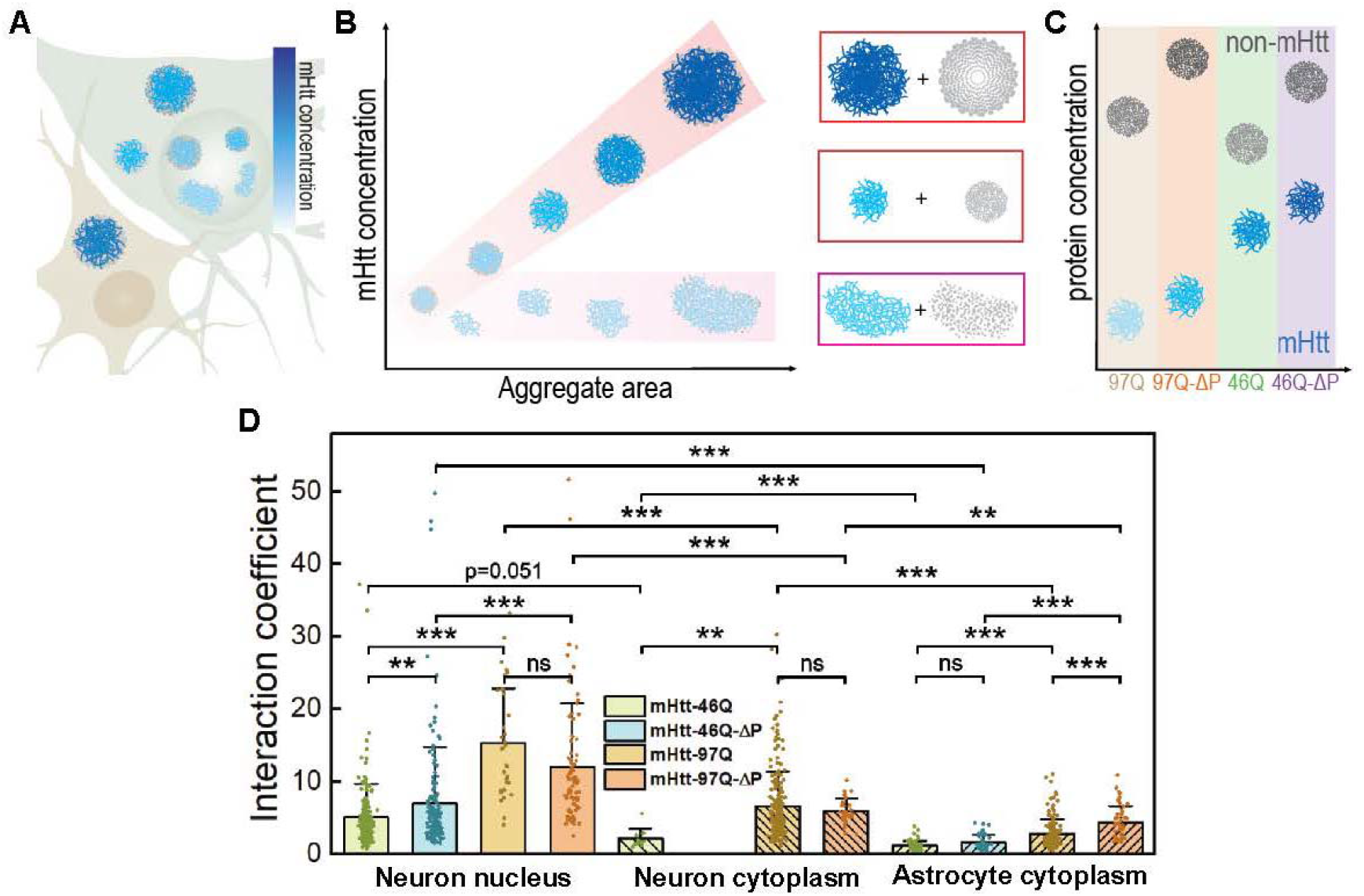
Proposed model of native neuronal polyQ aggregates. (A) Subcellular distribution of polyQ aggregates in neurons and astrocytes. (B) Our proposed quantitative model for polyQ aggregate formation and growth, pink: gel-like aggregates; coral: solid-like aggregates. Right panel shows the spatial distribution of mHtt proteins (Blue) and non-mHtt proteins (Gray) within corresponding different types of polyQ aggregates. (C) Comparison of c(mHtt, blue) and c(non-mHtt, gray) for polyQ aggregates across different mHtt constructs, including mHtt-97Q, mHtt-46Q, mHtt-97Q-ΔP, and mHtt-46Q-ΔP. (D) Bar diagram of proposed interaction coefficients (ICs, defined as c(non-mHtt)/c(mHtt))) in different cell types, subcellular compartments across different mHtt constructs, including mHtt-97Q, mHtt-46Q, mHtt-97Q-ΔP, and mHtt-46Q-ΔP. ns: not significant, * p<0.05, ** p < 0.01, *** p < 0.001.

The strong linear correlation between c(mHtt) and aggregate sizes enables precise quantification of aggregate growth. We observed that the slopes in the fitting curves remained consistent in both neurons and astrocytes, suggesting a similar growth trend across different cell types. However, the intercepts vary, which points to different critical concentrations required for aggregation initiation and likely reflects the heightened susceptibility to mHtt perturbations in neurons compared to astrocytes. Notably, we found that c(mHtt) in mHtt-46Q aggregates is approximately twice that of similarly sized mHtt-97Q aggregates, resulting in a two-fold steeper slope in the mHtt-46Q fitting curve (Fig. S8C). This suggests that mHtt-46Q has a weaker aggregation ability with a higher c(mHtt) requirement for aggregation, consistent with the fact that longer polyQ repeats result in higher aggregation propensity and earlier age of disease onset for HD.(41, 57) Interestingly, the Q count in mHtt-97Q is about twice that of mHtt-46Q, which raise the possibility that aggregation might be driven by overall Q concentrations which are the same between mHtt-46Q and mHtt-97Q aggregates rather than c(mHtt).

One major debate for poly-Q research is whether the formation of polyQ aggregates is harmful or protective. Recent evidence suggests that soluble mHtt oligomers interact more extensively with cellular proteins compared to mHtt in polyQ aggregates, making soluble oligomers more toxic. (5, 9, 11-14) However, previous experiments have struggled to accurately measure the total amount of non-mHtt proteins increasing within the aggregates due to the need for extensive polyQ extraction. Our platform offers a live-cell solution to this challenge. We hence introduced an interaction coefficient (IC), defined as the concentration ratios of non-mHtt over mHtt proteins (Fig. 6C). This number quantitatively represents the “dragging” potential of each mHtt protein in pulling multiple copies of non-mHtt structural or functional proteins (Fig. 6D), which may serve as a quantitative indicator of mHtt toxicity for its impact on cellular function.

ICs are higher in neurons compared to those in astrocytes, consistent with the notion that mHtt exerts higher toxicity in neurons than astrocytes (Fig. 6D). Interestingly, ICs are much higher for nuclear aggregates than for cytoplasmic aggregates, further supporting the hypothesis that nuclear aggregates are more toxic, as demonstrated previously by cellular function studies.(22, 59, 60) As mentioned in the introduction, younger patients tend to exhibit nuclear aggregates, while older patients more commonly show cytoplasmic aggregates. Our findings highlight the distinct difference between nuclear and cytoplasmic aggregates, suggesting a potential pathway to better understand how the subcellular localization of polyQ aggregates may impact HD progression. In addition, as size increases, ICs decrease for most aggregates other than the gel-like ones (Fig. 6D), suggesting that the growth of aggregates may help mitigate cellular functional loss, supporting the protective role of aggregate formation. Furthermore, ICs of mHtt-46Q-ΔP and mHtt-97Q-ΔP aggregates are higher than those of mHtt-46Q and mHtt-97Q (Fig. 6D), consistent with increased toxicity after deleting the proline-rich region. All this evidence points to the potential effectiveness of ICs as a new quantitative indicator of aggregate toxicity, which may find utility in high-throughput screening of drug discovery for HD. Interestingly, we observed that gel-like nuclear aggregates exhibited significantly higher IC values compared to solid-like nuclear aggregates (Fig. S9B), suggesting potential differences in the toxicity associated with these distinct aggregation types. Of course, more functional verifications are needed to confirm the role of IC, which is an important direction for our future work. Multidimensional information from tandem mass spectrometry with q-aggSRS should further confirm the protein identity in the non-mHtt protein pool.

## Materials and Methods

### Primary neuron culture

Primary rat hippocampal neurons were isolated from neonatal Sprague–Dawley rat (CD (Sprague– Dawley) IGS rat, Charles River) pups with a protocol (IA22-1835) approved by Caltech’s Institutional Animal Care and Use Committee (IACUC). The brains were dissected from the skull and placed into a 10-cm Petri dish with ice-chilled Hanks’ balanced salt solution (Gibco). The hippocampus was isolated from the brains under a dissection scope, cut into small pieces (∼0.5□mm), and incubated with 5□ml of Trypsin-EDTA (0.25%, Gibco) at 37□°C with 5% CO_2_ for 15□min. The Trypsin-EDTA liquid was aspirated and replaced with 2□ml of DMEM containing 10% FBS to stop the digestion. The tissue fragments were moved into 2□ml of neuronal culture medium (Neurobasal A medium, B-27 supplement, 2 mM GlutaMAX supplement, Thermos Fisher, and 1× penicillin-streptomycin) and dispersed by repeated pipetting several times. The supernatant was collected and further diluted by neuronal culture medium to a final cell density of 9□×□104□cells□ml^−1^. A 0.7-ml volume of cell suspension was added to each well of a 24-well plate on coated 12-mm circular cover glass. For pre-coating, sterile 12-mm circular cover glass was incubated with 100□μg/ml poly-d-lysine (Sigma) solution at 37□°C with 5% CO_2_ for 24□h in a 24-well plate. The 12-mm circular cover glass was washed twice with ddH_2_O and incubated with 10□μg/ml laminin mouse protein (Gibco) solution at 37□°C with 5% CO_2_ overnight. Thereafter, the 12-mm circular cover glass was washed twice with ddH_2_O and allowed to dry at room temperature inside a biosafety cabinet. Half of the neuron culture medium was replaced with fresh medium every four days. All the medium changes were done after pre-heating to 37°C.

### AAV packaging

We used the pAAV backbone for building plasmid constructs. The inserted gene was a mutant HTT gene encoding the exon 1 of mHtt protein. The model mHtt protein contains an N-terminal 17 a.a. fragments (MATLEKLMKAFESLKSF), the polyQ tract (46Q or 97Q), and the proline-rich region PPPPPPPPPPPQLPQPPPQAQPLLPQP QPPPPPPPPPPGP). For mHtt protein without EGFP, the C-terminal is AVAEEPL HRP*, while for mHtt-EGFP protein, there is a linker (AVAEEPLHRPGSSPVAT) between mHtt and EGFP (MVSKGEELFTGVVPILVELDGDVNGHKFSVSGEGEGDATYGKLTLKFICTTGKLPVPWPTLVTTLTYGVQCFSRYPDHMKQHDFFKSAMPEGYVQERTIFFKDDGNYKTRAEVKFEGDTLVNRIELKGIDFKEDGNILGHKLEYNYNSHNVYIMADKQKNGIKVNFKIRHNIEDGSVQLADHYQQNTPIGDGPVLLPDNHYLSTQSALSKDPNEKRDHMVLLEFVTAAGITLGMDELYK*). The expression of mHtt protein was initiated by either a general promoter, CMV (cytomegalovirus) promoter or a neuron-specific promoter hSYN promoter (Human synapsin 1 gene promoter). The plasmids were sent to the CLOVER center at Caltech to produce AAV-DJ vectors. The typical titer is around 1E+13 genome copies (GCs)/mL with a normal yield of AAV vectors.

### AAV transduction and medium change

AAV vectors were added to neurons at DIV 6-8 for about 5E9 GCs per well for the incubation of about 24 hours. A full-medium change from the normal medium to the medium with Gln-d_5_ or Gln-d_5_+Val-d_8_ was performed after the AAV incubation, and cells were incubated with the new medium for 3-5 days. Gln-d_5_ neuron culture medium was made by replacing GlutaMAX with 2 mM Gln-d_5_ (Cambridge Isotope Laboratories) in Neurobasal A medium which is glutamine-deficient. Gln-d_5_ and Val-d_8_ complete neuron culture medium was made from an amino-acid-deficient Neurobasal powder (BioTrend) with adding other normal amino acids and Gln-d_5_ & Val-d_8_ (Cambridge Isotope Laboratories) corresponding to the concentrations listed in Neurobasal A medium formula. For the pulse-chase experiment, the full-medium change from the normal medium to the medium with Gln-d_5_ was done after the first round of SRS imaging. Cells were incubated with the new medium for an additional 8 h before the second round of SRS imaging. All the medium changes were done after pre-heating to 37°C.

### Stimulated Raman Scattering (SRS) Microscopy

A picoEmerald laser system (Applied Physics and Electronics) was used as the light source for SRS microscopy. Briefly, it produces 2 ps pump (tunable from 770 nm – 990 nm, bandwidth 0.7 nm, spectral bandwidth ∼ 10 cm^-1^) and Stokes (1032 nm, spectral bandwidth 10 cm^-1^) pulses with 80 MHz repetition rate. Stokes beam is modulated at 20 MHz by an internal electro-optic modulator. The spatially and temporally overlapped Pump and Stokes beams are introduced into an inverted laser-scanning microscope (FV3000, Olympus), and then focused onto the sample by a 25X water objective (XLPLN25XWMP, 1.05 N.A., Olympus). Transmitted Pump and Stokes beams are collected by a high N.A. condenser lens (oil immersion, 1.4 N.A., Olympus) and pass through a bandpass filter (893/209 BrightLine, 25mm, Semrock) to filter out Stokes beam. A large area (10×10 mm) Si photodiode (S3590-09, Hamamatsu) is used to measure the pump beam intensity. A 64 V reverse-biased DC voltage is applied to the photodiode to increase the saturation threshold and reduce response time. The output current is terminated by a 50Ω terminator and pre-filtered by a 19.2-23.6-MHz band-pass filter (BBP-21.4+, MiniCircuits) to reduce laser and scanning noise. The signal is then demodulated by a lock-in amplifier (SR844, Stanford Research Systems) at the modulation frequency. The in-phase X output is fed back to the Olympus IO interface box (FV30-ANALOG) of the microscope. 30 µs time constant is set for the lock-in amplifier. Correspondingly, 80 µs pixel dwell time is used, which gives a speed of 21.3 s/frame for a 512-by-512-pixel image, with two frame-averaging. Laser powers are monitored throughout image acquisition by an internal power meter and power fluctuations are controlled within 1%. 16-bit greyscale images were acquired by Olympus Fluoview 3000 software.

Right before imaging, the cover glass with neurons was taken out to make an imaging chamber (Grace Bio-Labs SecureSeal™ imaging spacer, diam. × thickness 9 mm × 0.12 mm) filled with the original medium for SRS imaging. For the channel of CH_3_, CH_2_, CD on (for Gln-d_5_), CD off (for Gln-d_5_), CD on (for Val-d_8_), CD off (for unmixing), the wavelengths of pump lasers for SRS are 791.8 nm, 797.8 nm, 843.6 nm, 852.8 nm, 850.5 nm, 855 nm. For hyperspectral measurements, the wavelengths of the pump laser were tuned from 835 nm to 855 nm with a 0.5 nm interval. All images were processed using ImageJ software. Corresponding off-resonance images were subtracted.

### Materials and sample preparation for VISTA

Sodium acrylate (SA), N,N′-methylenebisacrylamide (BIS, 2%), ammonium persulfate (APS), tetramethylethylenediamine (TEMED), and sodium dodecyl sulfate (SDS) were purchased from Sigma-Aldrich. Acrylamide (AA) was purchased from Ambeed, Inc. The 1.0 M Tris solution was obtained from Biosolve, and nuclease-free water was from Ambion. The expansion protocol follows our previously published methodology. Stock solutions were prepared and stored under the following conditions: an incubation solution containing 30% AA in 4% paraformaldehyde (PFA) in 1× phosphate-buffered saline (PBS) was stored at 4°C. A gelation solution consisting of 7% SA, 20% AA, and 0.1% BIS in 1× PBS was stored at −20°C. APS and TEMED were dissolved in nuclease-free water to create a 10% (w/w) solution and stored at −20°C.

For hydrogel embedding, cell samples were incubated in a 30% AA and 4% PFA solution without prior fixation. This incubation was carried out for 16 hours at 37°C, followed by a wash with gelation solution at 4°C. The gelation solution, APS, and TEMED were thawed and kept at 4°C until use. Coverslips containing cells were placed at the bottom of a 4°C gelation chamber with the cell side facing up. A mixture of 90% gelation solution, 5% APS solution, and 5% TEMED solution was prepared and immediately pipetted onto the sample to ensure complete immersion before covering it with a glass slide. The chamber was maintained at 4°C for 1 minute before being transferred to a humidified incubator for 1 hour at 37°C.

Post-gelation, the coverslips with gels were incubated in a denaturing buffer (200 mM SDS, 200 mM NaCl, and 50 mM Tris in nuclease-free water, pH 8) in Petri dishes for 15 minutes at room temperature. Subsequently, they were transferred to 2 ml Eppendorf centrifuge tubes and heated in 1.5 ml of denaturing buffer at 95°C for 30 minutes. Initial expansions were performed immediately after denaturation at room temperature in water, with the water changed every hour for three times. The expanded gel was then shrunk in PBS overnight in the dark at 4°C. Under these conditions, the cells underwent a 2-fold linear isotropic expansion.

### Statistical analysis

All the statistical analyses were done using OriginPro 2024b software. All data are shown as means ± SEM. One-way analyses of variance (ANOVAs) followed by Tukey’s multiple comparisons test were used for single-variable comparisons. The statistical significance level was defined as: ns not significant, * p<0.05, ** p < 0.01, *** p < 0.001.

## Supporting Information

Detailed experimental procedures, derivation of absolute concentrations, and supplementary figures

## Acknowledgments

We acknowledge the Caltech CLOVER center, Institutional Animal Care and Use Committee (IACUC), and the Office of Laboratory Animal Resources (OLAR) for research resources. We thank Prof. Dennis A. Dougherty, Prof. Viviana Gradinaru, Prof. Shu-ou Shan, Charlene Kim, Dr. Shuzhang Liu, Dr. Jiajun Du, Rahuljeet (RJ) Chadha, Adrian Colazo, Noor Naji, and Berea Suen for fruitful discussions. This work is supported by an Alfred P. Sloan Research Fellowship (L.W.). X.B. acknowledges the support received from the Biotechnology Leadership Pre-Doctoral Training Program (BLP) in the Donna and Benjamin M. Rosen Bioengineering Center and the Barbara J. Burger Graduate Fellowship at Caltech. L.W. is a Heritage Principal Investigator supported by the Heritage Medical Research Institute.

## Author Contributions

L.W. supervised this project. X.B. and L.W. conceived and designed the research. X.B. performed the experiments and analyzed the data. L.L. helped with the VISTA experiment. K.M. helped with the experimental materials. The manuscript was written by X.B. and L.W., with input from all authors.

