## Supporting Information for "GFP-Free Live-Neuron Quantitative Imaging Reveals Compartmentalization and Growth Dynamics of PolyQ Aggregates"

^2^Current address: Department of Pharmaceutical Chemistry, University of California San Francisco, San Francisco, California 94115, USA

*Corresponding author: Lu Wei.

**Supporting Information Text**

**Identification of live neurons and astrocytes for aggregate analysis**

To image aggregates specifically in live neurons and astrocytes, we adopted Neuron-NeuO staining in live neuronal co-cultures for experimental analysis. NeuroFluor™ NeuO (STEMCELL) is a green membrane-permeable fluorescent probe that selectively labels primary and pluripotent stem cell-derived neurons in the presence of other brain cells.(1) The neurons were incubated with 0.25 μM NeuO for 1 hour and then were changed back to normal medium without NeuO for another 2 hours. In this case, neurons present clear green fluorescence with corresponding images in the CH_3_ SRS channel; astrocytes present as cells in the CH_3_ SRS channel but have no green fluorescence contrast (Fig. S3C).

We further benchmarked the quantitative results with other cell distinguishing strategies and found good consistency (Fig. S5A). First, we used the neuron-specific promoter, hSYN. Second, we added 2 uM 1-β-D-arabinofuranosylcytosine (Ara-C) to the culture medium at DIV 3 to obtain pure neuron cultures without astrocytes (major species other than neurons) or oligodendrocytes (minor species) (Fig. S3A). To specifically image aggregates in astrocytes, we also performed pure astrocyte cultures (Fig S3B), which were obtained by using Basal Eagle Media containing 10% fetal bovine serum (FBS, Invitrogen) and GlutaMAX supplement.

**Calculation of mHtt/non-mHtt concentrations**

We use 10 mM Gln-d_5_ solution as a standard to calibrate the SRS signal of CD bonds to the concentration of CD bonds (Fig. S1). We confirmed that SRS signals of CD bonds have a linear relationship with Gln-d_5_ concentration, and the CH Raman cross-section is the same as CD (confirmed by DMSO-d_6_ and DMSO comparison). All signals are processed with power normalization.

For non-mHtt proteins, the average length of proteins in eukaryotes is 438 amino acids.(2) On average there are 6 C-H bonds of each amino acid, with the consideration of relative abundance. To calculate the concentration of mHtt proteins (c(mHtt)) and non-mHtt proteins (c(non-mHtt)) within the same aggregates from measured CD signals and the CH/CD ratios, we can write the following two Equations (1) and (2):

$c\left( CD \right)=5*c\left( mHtt \right)*n\left( Q \right)+5*c\left( non-mHtt \right)*438*X\%$	(1)

$c\left( CH \right)=6*c\left( mHtt \right)*n\left( non-Q a.a. \right)+6*c\left( non-mHtt \right)*438*(1-X\%)$ (2)

Here $X\%$ indicates the incorporation ratio for Gln-d_5_ in the regular proteome of non-Htt proteins. We determine this number experimentally by incubating neurons with Gln-d_5_ medium without AAV transfection. After 0-6 days’ incubation, we took SRS images at both CH_3_ and CD channels to determine $X\%$. We found that the CH/CD ratios plateau from day 3 to day 6, shown below in Fig. S4, with an average CH/CD ratio of 55. We confirmed through d_7_-glucose labeling that *de novo* glutamine synthesis is negligible. So we have the following equation for $X\%$ as:

$$\frac{6*\left( 1-X\% \right)}{5*X\%}=55$$

Yielding $X\%$ of 2.1% after 3-6 days of Gln-d_5_ incubation, consistent with the number that Gln-d_5_ makes up 4.2% of the overall proteome.(3) We use 2.1% for all calculations in Eq (1) and Eq (2) for c(mHtt) and c(non-mHtt).

For different mHtt-exon1 constructs, $n\left( Q \right)$, the number of glutamines, and $n\left( non-Q a.a. \right)$, the number of non-glutamine amino acids, are exactly known and shown in the table below in Table S1. Taking these numbers and each measured CH/CD ratio back into Eq. (1) and (2) can solve for c(mHtt) and c(non-mHtt), corresponding to the mHtt concentration and non-mHtt cytosolic protein concentrations within the same aggregates.

**Table S1: Number of glutamine (Q) and non-glutamine amino acid (non-Q a.a.) in each mHtt construct sequence.**

|  | 97Q | 97Q-$\Delta$P | 97Q-EGFP | 46Q | 46Q-$\Delta$P | 97Q-EGFP |
| --- | --- | --- | --- | --- | --- | --- |
| $n\left( Q \right)$ | 103 | 97 | 111 | 52 | 46 | 60 |
| $n\left( non-Q a.a. \right)$ | 61 | 27 | 299 | 61 | 27 | 299 |

**Unmixing for two-color imaging of Gln-d_5_ and Val-d­_8_**

Two images were taken at the channel of 2164 cm^-1^ (pump laser 843.6 nm) and 2068 cm^-1^ (pump laser 850.5 nm). Background at the channel of 2006 cm^-1^ (pump laser 855 nm) was subtracted before further analysis. According to the spectra of aggregates incorporated with Gln-d_5_ and Val-d_8_ (Fig 3B, bottom), we can obtain two linear equations below.

$${SRS}_{2164}={SRS}_{{Gln-d}_{5}}+0.6*{SRS}_{{Val-d}_{8}}$$

$${SRS}_{2068}=0.1*{SRS}_{{Gln-d}_{5}}+{SRS}_{{Val-d}_{8}}$$

So we can obtain the equations for unmixing to get pure signals from Gln-d_5_ and Val-d_8_ labeling channels as:

$${SRS}_{{Gln-d}_{5}}=\frac{10*{SRS}_{2164}-6*{SRS}_{2068}}{9.4}$$

$${SRS}_{{Val-d}_{8}}=\frac{10*{SRS}_{2068}-{SRS}_{2164}}{9.4}$$

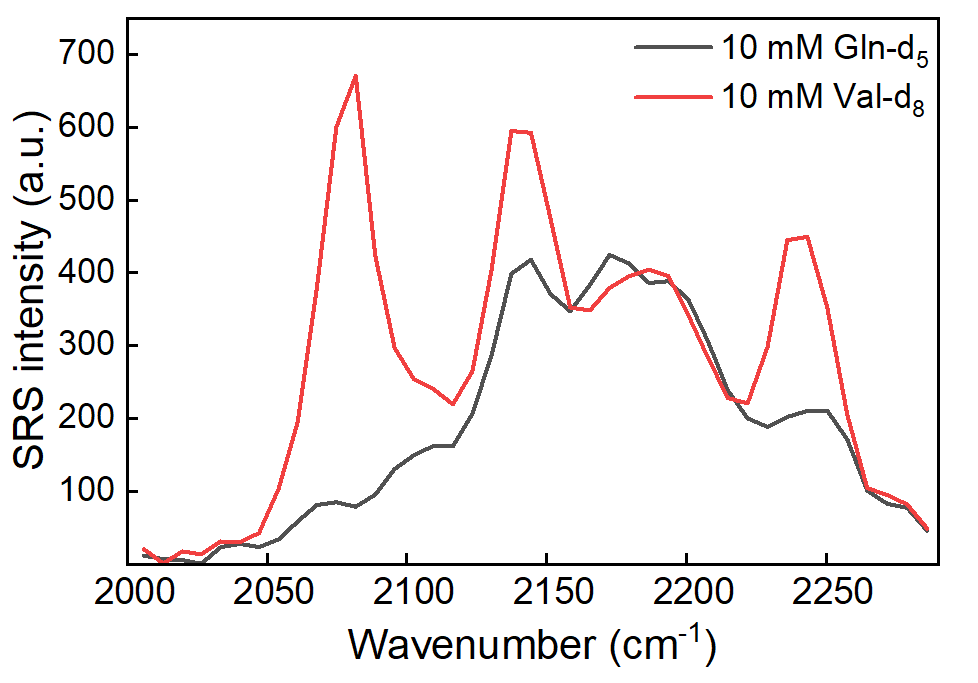

**Fig S1. SRS spectra of 10 mM Gln-d_5_ and Val-d_8_ in water solutions.**

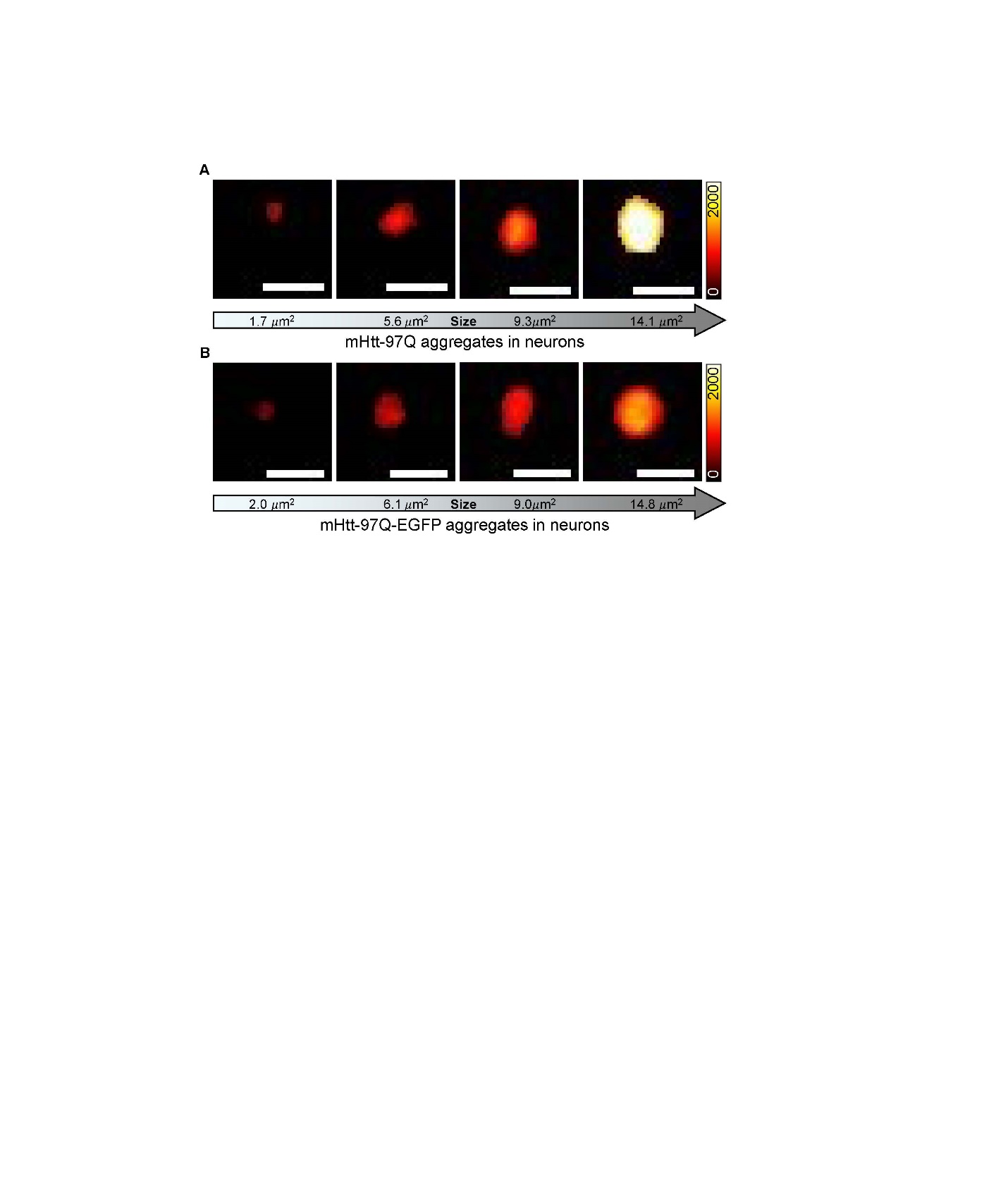

**Fig S2. Comparison of SRS CD intensity of polyQ aggregates in neurons for mHtt-97Q (top, A) or mHtt-97Q-EGFP (bottom, B) sequences.** (A) SRS images at the CD on-off channel (on, 2164 cm^-1^; off, 2036 cm^-1^) for 4 representative mHtt-97Q aggregates with increasing sizes in neurons. (B) SRS images at the CD on-off channel (on, 2164 cm^-1^; off, 2036 cm^-1^) for 4 representative mHtt-97Q-EGFP aggregates of close sizes to that shown in (a) in neurons. Scale bar: 5μm. Color bars are kept the same for (A) and (B).

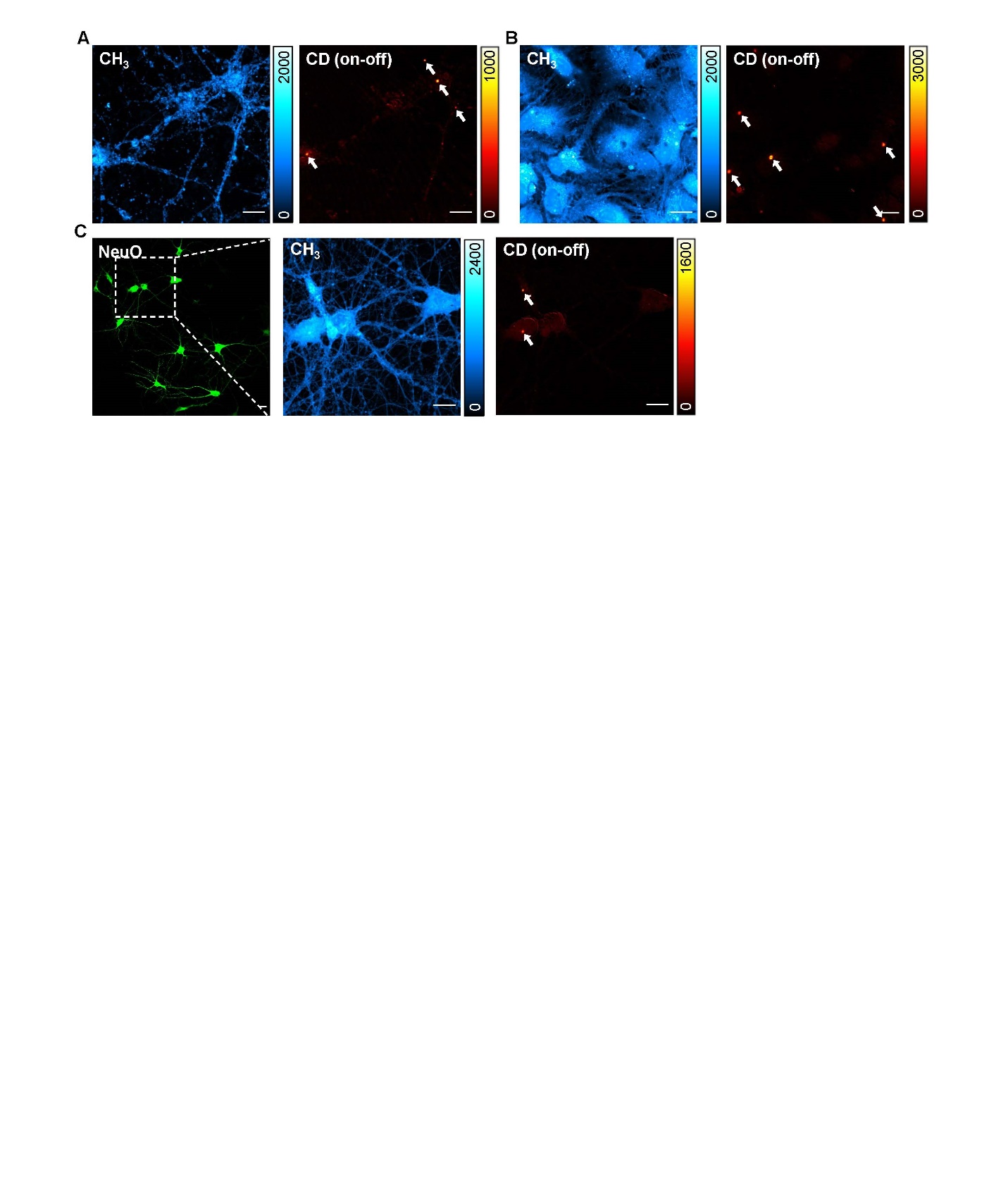

**Fig S3. Q-aggSRS images of polyQ aggregates in neurons and astrocytes with different cell-distinguishing strategies.** (A) SRS images at the CH_3_ channel (left, 2940 cm^-1^) and CD on-off channel (right, on, 2164 cm^-1^; off, 2036 cm^-1^) in the same set of pure neuron culture containing mHtt-97Q aggregates (arrowed in the CD image). (B) SRS images at the CH_3_ channel (left, 2940 cm^-1^) and CD on-off channel (right, on, 2164 cm^-1^; off, 2036 cm^-1^) in the same set of pure astrocyte culture containing mHtt-97Q aggregates (arrowed in the CD image). (C) Fluorescence image (left, NeuO) in live neuronal co-cultures containing mHtt-97Q aggregates with NeuO staining. Corresponding SRS images at the CH_3_ channel (middle, 2940 cm^-1^) and CD on-off channel (right, on, 2164 cm^-1^; off, 2036 cm^-1^) at the zoomed-in region in fluorescence (white-dashed box) are shown. mHtt-97Q aggregates are arrow-indicated in the CD image, indicating the top one in the astrocyte and the bottom one in the neuron. Scale bar: 20 μm.

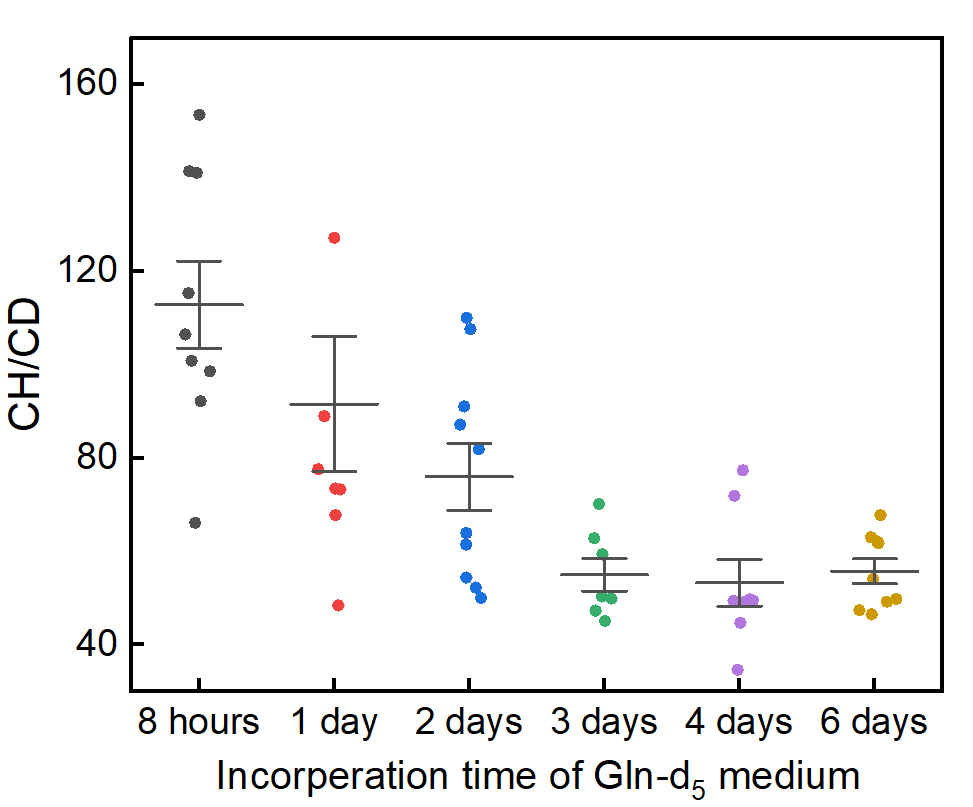

**Fig. S4 CH/CD ratios dependence on Gln-d_5_ medium incubation time without transductions of AAV packed mHtt plasmids.**

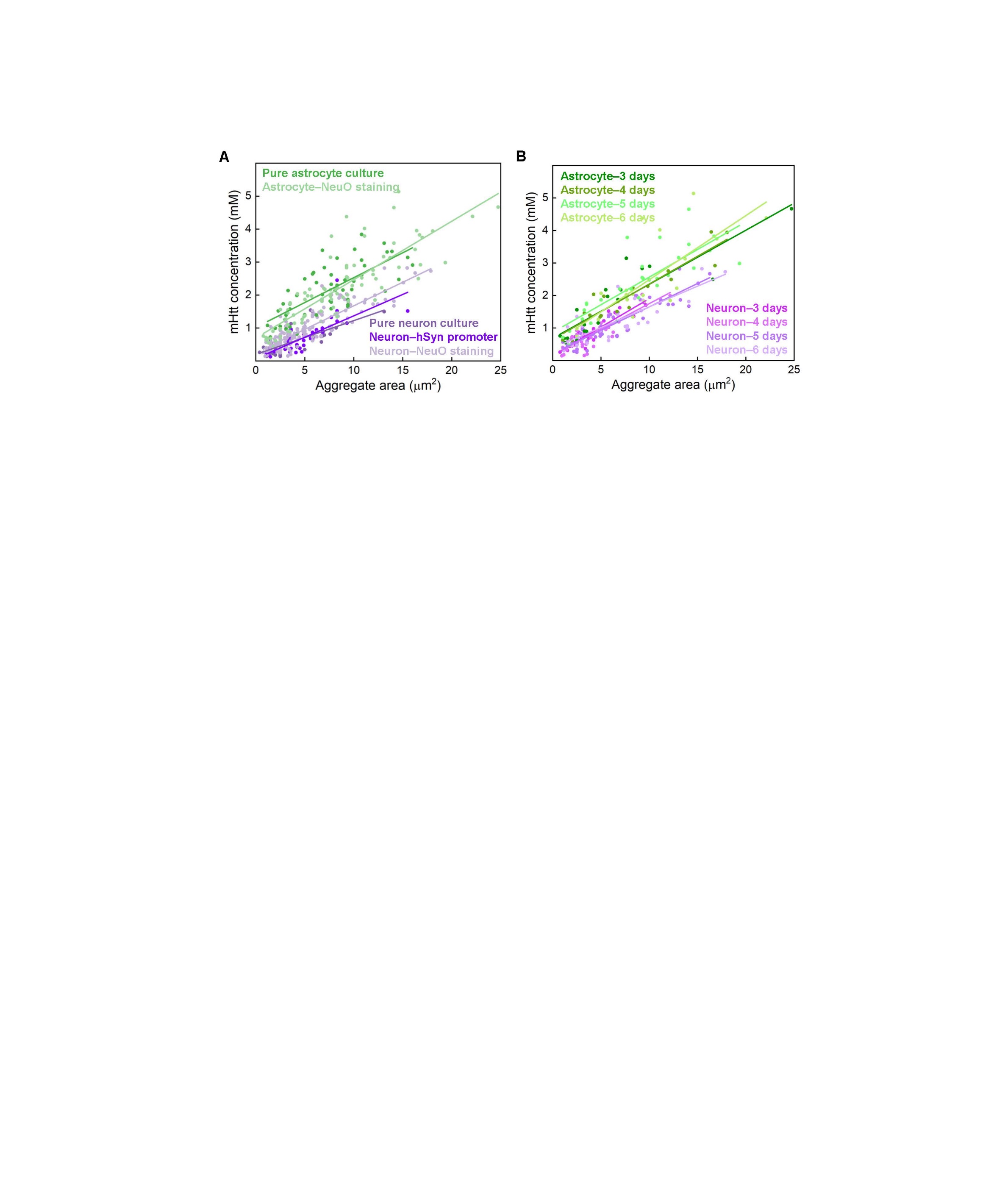

**Fig S5. Demonstration of validity of calculated mHtt concentration (c(mHtt)) with minimal influence from cell-differentiation methods and expression time.** (A) Scatter plots with linear fittings between c(mHtt) and aggregate areas of polyQ aggregates in neurons and astrocytes by different cell-differentiation methods mentioned above. Pure neuron culture: $y=0.20x+0.10$, Pearson’s $r=0.74$; Neuron-hSyn promoter: $y=0.13x+0.074$, Pearson’s $r=0.78$; Neuron-NeuO staining: $y=0.14x+0.25$, Pearson’s $r=0.88$; Pure astrocyte culture: $y=0.15x+1.02$, Pearson’s $r=0.74$; Neuron-NeuO staining: $y=0.18x+0.70$, Pearson’s $r=0.84$. (B) Scatter plots with linear fittings between c(mHtt) and aggregate areas of polyQ aggregates in neurons and astrocytes in different post-AAV incubation times. Neuron-3 days: $y=0.18x+0.15$, Pearson’s $r=0.93$; Neuron-4 days: $y=0.16x+0.18$, Pearson’s $r=0.82$; Neuron-5 days: $y=0.14x+0.22$, Pearson’s $r=0.90$; Neuron-6 days: $y=0.13x+0.38$, Pearson’s $r=0.81$; Astrocyte-3 days: $y=0.17x+0.68$, Pearson’s $r=0.88$; Astrocyte-4 days: $y=0.17x+0.65$, Pearson’s $r=0.94$; Astrocyte-5 days: $y=0.17x+0.86$, Pearson’s $r=0.82$; Astrocyte-6 days: $y=0.20x+0.51$, Pearson’s $r=0.75$;

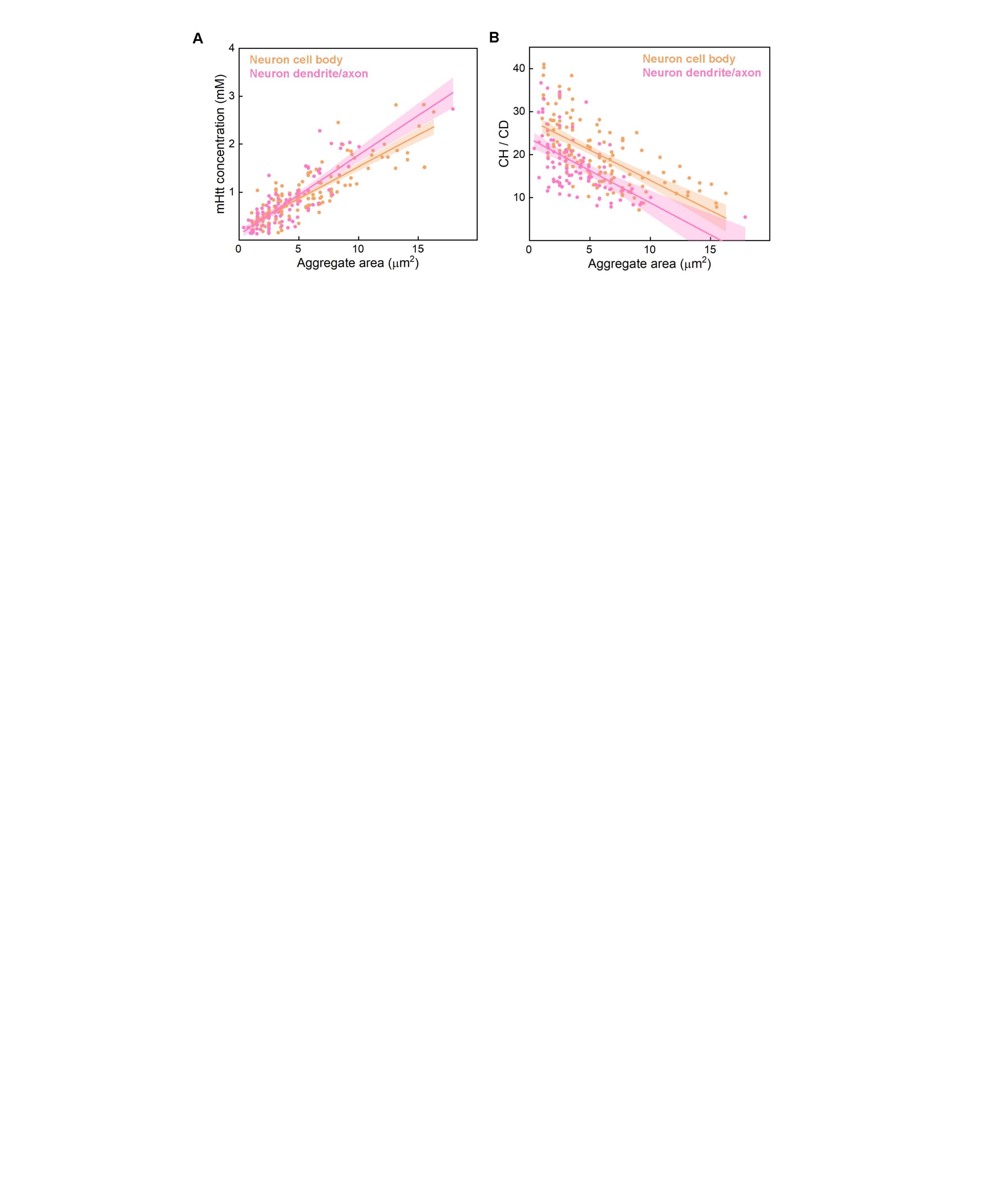

**Fig S6. Quantitative analysis of mHtt-97Q aggregates in neuronal subcellular localizations of the cell body and dendrite/axon.** (A) Scatter plots with linear fittings between c(mHtt) and aggregate areas of polyQ aggregates in the neuron cell body (orange, $y=0.13x+0.20$, Pearson’s $r=0.84$) and dendrite/axon (pink, $y=0.17x+0.11$, Pearson’s $r=0.84$). (B) Scatter plots with linear fittings between CH/CD ratios and aggregate areas of polyQ aggregates in the neuron cell body (orange, $y=-1.41x+28.12$, Pearson’s $r=-0.65$) and dendrite/axon (pink, $y=-1.50x+23.78$, Pearson’s $r=-0.58$). A 95% confidence band is plotted with the fitted line.

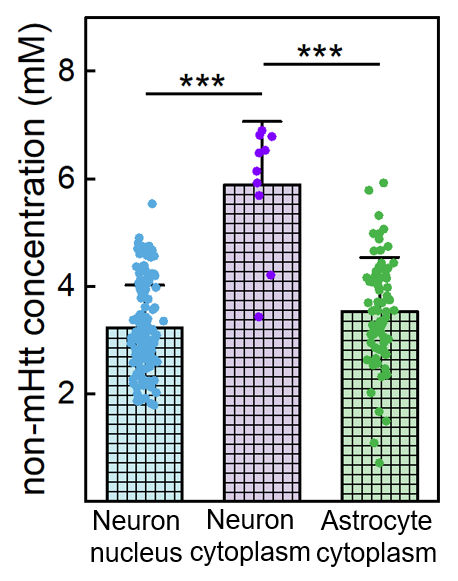

**Fig S7. Non-mHtt concentrations of mHtt-46Q aggregates in different subcellular compartments, including neuron nucleus (blue), neuron cytoplasm (purple), and astrocyte cytoplasm (green), *** p < 0.001. No plot is shown for astrocyte nucleus due to almost 0% aggregate appearance.**

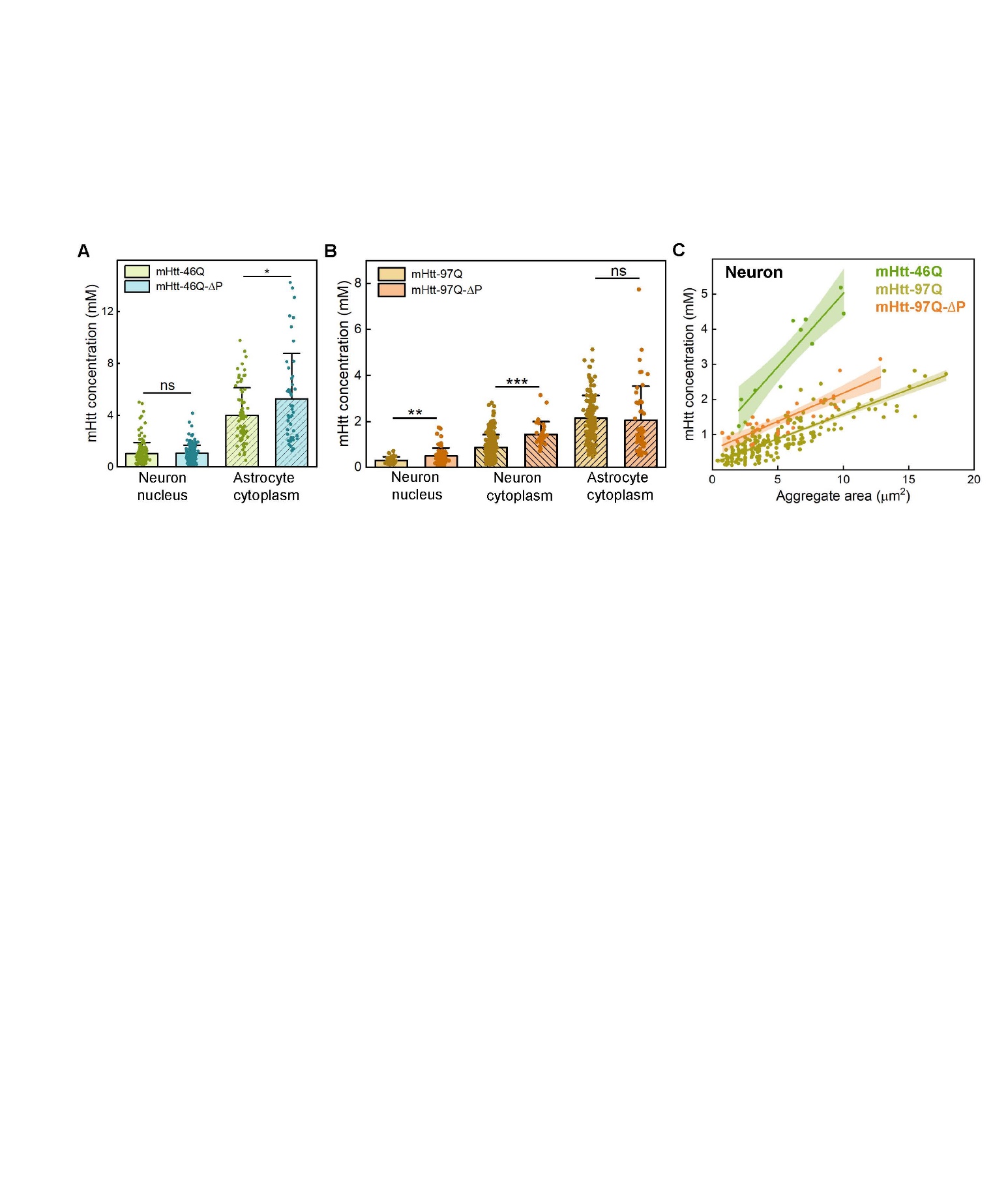

**Fig S8. Quantitative measurement of polyQ aggregates with different mHtt constructs in neurons and astrocytes.** (A-B) c(mHtt) of polyQ aggregates in different subcellular compartments, including neuron nucleus, neuron cytoplasm, and astrocyte cytoplasm, for mHtt-46Q/mHtt-46Q-ΔP (A) and mHtt-97Q/mHtt-97Q-ΔP (B), ns: not significant, * p < 0.05, ** p < 0.01, *** p < 0.001. No neuronal cytoplasm data is shown in (A) due to an insufficient number of aggregates detected, as the majority of aggregates are in neuronal nuclei. (C) Scatter plots with linear fittings between c(mHtt) and aggregate areas of cytoplasmic aggregates in neurons. mHtt-46Q: $y=0.42x+0.84$, Pearson’s $r=0.92$; mHtt-97Q: $y=0.14x+0.18$, Pearson’s $r=0.84$; mHtt-97Q-$\Delta$P: $y=0.16x+0.57$, Pearson’s $r=0.84$.

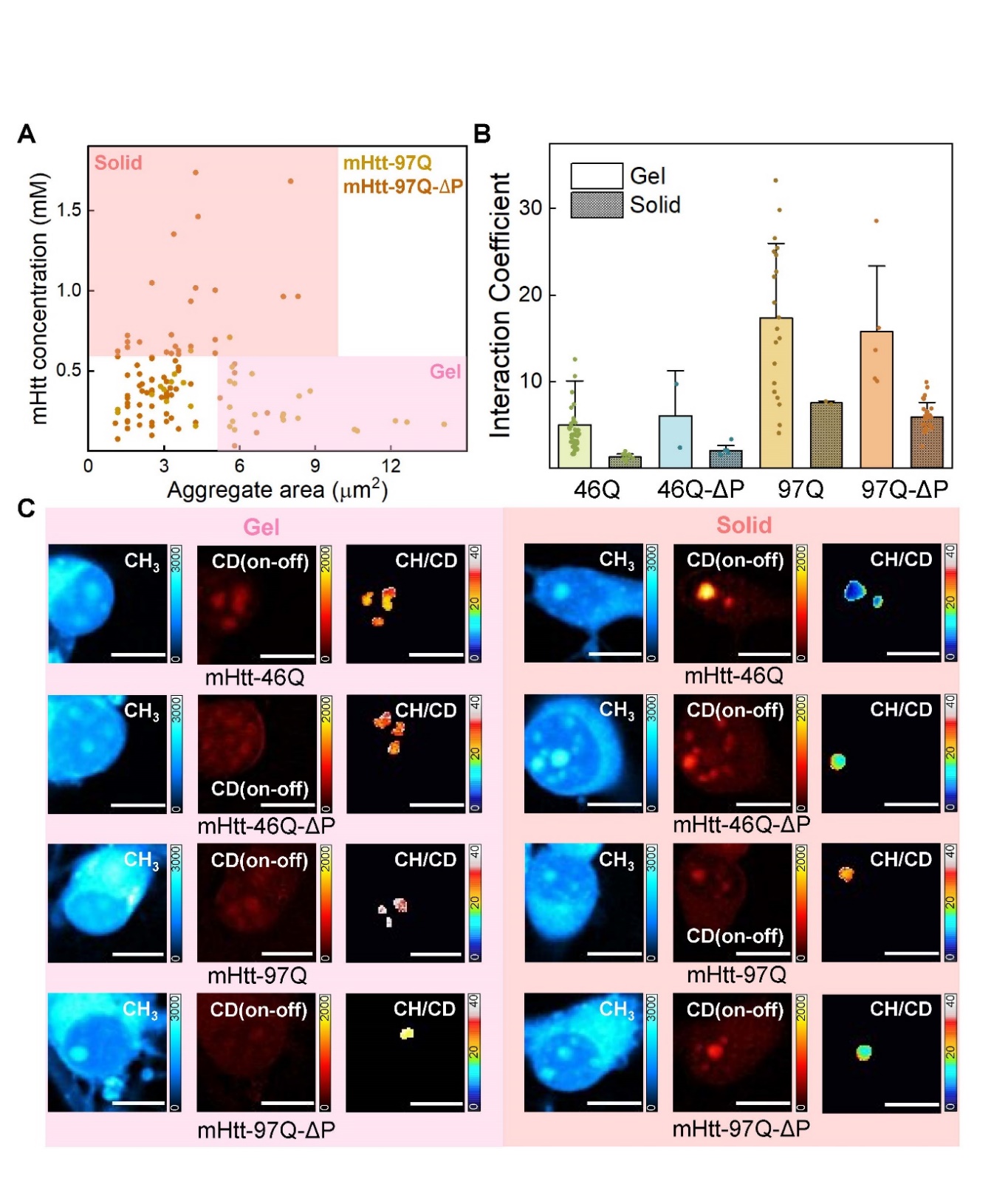

**Fig S9. Investigation of gel-like and solid-like nuclear aggregates with different mHtt constructs.** (A) Scatter plots of c(mHtt) vs aggregate areas for mHtt-97Q and mHtt-97Q-ΔP, indicating a bifurcation of concentration-area dependence. The pink region (lower right quadrant) indicates the nuclear gel-like aggregates, which feature low concentration-area dependence; the coral region (upper left quadrant) indicates the nuclear solid-like aggregates, which feature high concentration-area dependence. (B) Bar diagram of interaction coefficients (ICs, c(non-mHtt)/c(mHtt))) of gel-like (solid-fill) and solid-like (diagonal-line patterned) nuclear aggregates for different mHtt constructs of mHtt-46Q, mHtt-46Q-ΔP, mHtt-97Q, and mHtt-97Q-ΔP. (C) Q-aggSRS images of gel-like (left panel, pink) and solid-like (right panel, coral) nuclear aggregates across constructs of mHtt-46Q, mHtt-46Q-ΔP, mHtt-97Q, and mHtt-97Q-ΔP. For each set of three images: left: the CH_3_ channel (2940 cm^-1^), middle: the CD on-off channel (on, 2164 cm^-1^; off, 2036 cm^-1^), right: CH/CD ratios of the masked representative gel-like (left panel) or solid-like (right panel) aggregates. scale bar: 10 μm.
